# Nrl is dispensable for specification of rod photoreceptors in adult zebrafish contrasting a deeply conserved requirement earlier in ontogeny

**DOI:** 10.1101/2020.06.26.173930

**Authors:** A. Phillip Oel, Gavin J. Neil, Emily M. Dong, Spencer D. Balay, Keon Collett, W. Ted Allison

## Abstract

The transcription factor NRL (Neural Retinal Leucine-zipper) has been canonized, appropriately enough, as the master regulator of photoreceptor cell fate in the retina. NRL is necessary and sufficient to specify rod cell fate and to preclude cone cell fate in mice. By engineering zebrafish we tested if NRL function has conserved roles beyond mammals or beyond nocturnal species, i.e. in a vertebrate possessing a greater and more typical diversity of cone sub-types. Here, transgenic expression of a Nrl homolog from zebrafish or mouse was sufficient to convert developing zebrafish cones into rod photoreceptors. Zebrafish *nrl*^-/-^ mutants lacked rods (and had excess UV-sensitive cones) as young larvae, thus the conservation of Nrl function between mice and zebrafish appears sound. These data inform hypotheses of photoreceptor evolution through the Nocturnal Bottleneck, suggesting that a capacity to favor nocturnal vision is a property of *NRL* that predates the emergence of early mammals. Strikingly, however, rods were abundant in adult *nrl*^-/-^ null mutant zebrafish. Rods developed in adults despite Nrl protein being undetectable. Therefore a yet-to-be-revealed non-canonical pathway independent of *nrl* is able to specify the fate of some rod photoreceptors.

**Highlights:**

- Nrl is conserved and sufficient to specify rod photoreceptors in zebrafish retina
- Nrl is necessary for rod photoreceptors in early ontogeny of zebrafish larvae
- Zebrafish Nrl is functionally conserved with mouse and human NRL
- Remarkably, Nrl is dispensable for rod specification in adult zebrafish

## INTRODUCTION

Rods and cones are the ciliary photoreceptors used by vertebrates to enable vision across a broad range of circumstances. Rod photoreceptors enable vision in dim conditions, while cone photoreceptors convey wavelength-specific information, enable high acuity and can operate in brightly lit environments. Retinas with both rods and cones are known as duplex retinas, and the basic features of the duplex retina are present even among some of the earliest branching vertebrates, the lampreys [1-3].

The visual photoreceptors are among the best-studied neurons with respect to developmental programs and gene regulatory networks. Photoreceptor precursor cells of the developing mouse retina are thoroughly studied, and an elegantly simple gene regulatory network determines all rod and cone cell fates. As the precursor cell exits its terminal mitosis, expression of the bZIP transcription factor NRL directs it to a rod fate (schematized in Fig. 1A); without NRL expression it develops as a cone [4-7]. With high activity of the thyroid hormone receptor THRB, the presumptive cone will develop into the medium (green) wavelength light-sensitive M-cone (the ancestral red cone, expressing LWS opsin). Without THRB activity, it becomes a short wavelength (UV/blue) light-sensitive S-cone (the ancestral UV cone expressing SWS1 opsin) [8]. This efficient two-factor specification model is expected to be sufficient to generate all photoreceptor diversity in most all eutherian mammals, which have lost the ancestral blue and green light-sensitive cone subtypes [9]. In typical non-mammalian vertebrates, that have four cone subtypes (and rods), the blue and green cone specification remains unexplained.

**Figure 1.**
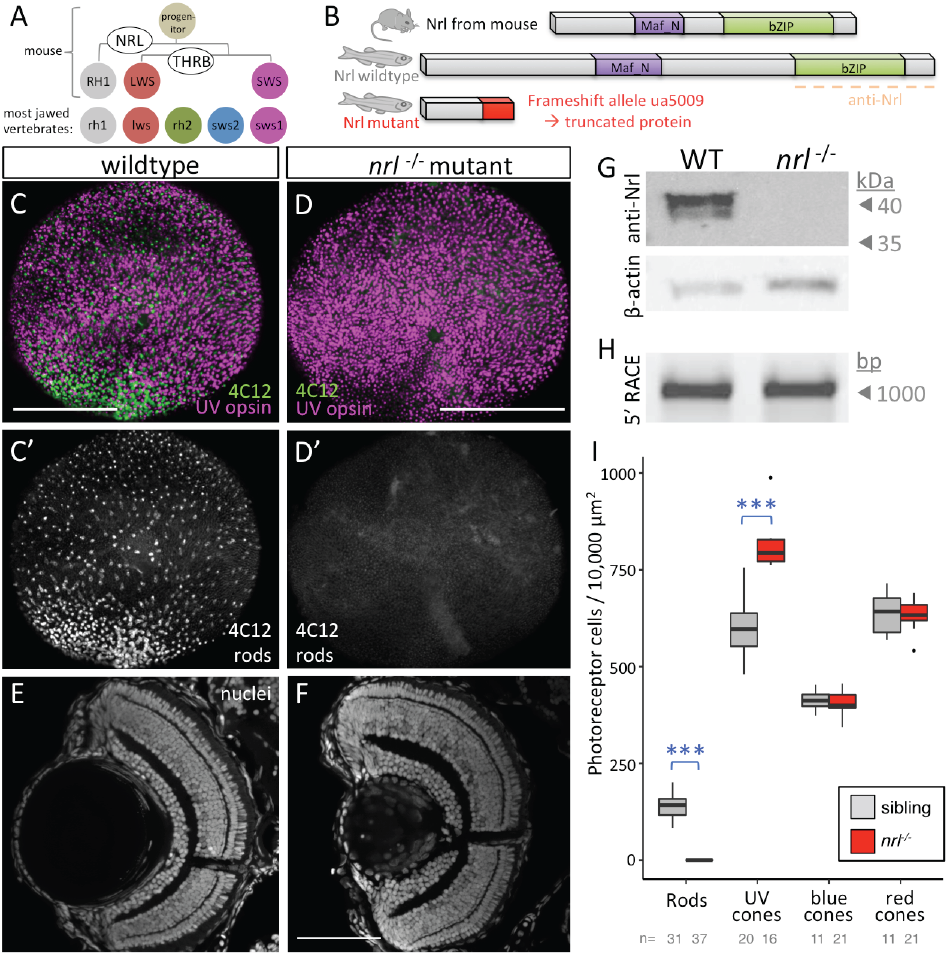
Nrl is conserved and required for rod specification in larval zebrafish. **A**. Nrl is the master regulator of photoreceptor specification in mice, being both necessary and sufficient for rod photoreceptor development from progenitor cells. Elegantly simple models can account for cell fate specification and generating the full complement of photoreceptor types in mice (and all mammals studied) using only two factors, NRL and THRB (thyroid hormone receptor β) to generate rods, red cones and blue cones expressing RH1, LWS and SWS opsins, respectively. However most vertebrates possess additional cone subtypes. **B**. Zebrafish Nrl protein is recognizably similar to mammalian homologs. An *nrl*^-/-^ null mutation was engineered via CRISPR that is predicted to truncate the protein (see also Supp Fig. S1). **C**,**D**. CRISPR-engineered null Nrl mutants lack rods in larval zebrafish, matching the phenotype of adult *NRL*^*-/-*^ knockout mice. Flat-mounted retina from larval zebrafish show that rods are absent in *nrl*^-/-^ larvae, as detected by rod immunomarker 4C12. The typical broad distribution of rods (B’) is absent (C’) and the decrease in rod cell abundance is consistent, nearly complete and robustly significant (**H**). **E**,**F**. Retina of *nrl*^-/-^ larvae is broadly normal in its lamination. **G**. Nrl protein is not detectable in *nrl*^-/-^ retina and antibody validation is supported by a doublet band (presumably reflecting SUMOylation as per mammalian homologs of Nrl) appearing at the predicted size of 44.2 kDa (See Fig. S1 for full immunoblots and quantification). **H**. Assessment of *nrl* transcript by 5’RACE (Random Amplification of cDNA Ends) does not reveal any unexpected transcripts such as those with cryptic exons that could be imagined to make functional protein. **I**. Quantification of photoreceptor types in *nrl*^-/-^ larvae confirms the consistent absence of rod cells, and a concomitant increase in UV cone abundance. Box and whisker plots first thru third quartile and distribution of data, respectively, after excluding outliers. ***p<0.001; bar in D is 100 um. Number of individual larvae examined (n) is indicated.

Thyroid hormone, and likely its receptor TRB2, control red cone development in chicken [10], and *trb2* regulates red cone versus UV cone fate in zebrafish [11-13], demonstrating substantial conservation of the photoreceptor specification program in tetrachromats. However, homologs of NRL have not been disrupted or manipulated sufficiently to appreciate its role(s) in species outside of mammals.

Previous work has shown *nrl* is expressed in or near the rod photoreceptors of zebrafish larvae [14, 15]. Xenopus embryos expressing lipofected Xenopus *Nrl* showed rhodopsin immunoreactivity in lipofected cells [16]. This provides tentative support for *nrl* playing a role in photoreceptor development outside of mammals. However, while mouse *NRL* is expressed detectably in developing lens tissue [17], lipofected human *NRL* did not promote lens fibre cell differentiation as Xenopus *Nrl* did [16], suggesting divergent activities for the gene homologs. Underscoring this, the avian lineage has lost any detectable ortholog of *NRL*, and likely relies on the *NRL*-family-member (a long Maf) gene *MAFA* for rod specification [18]; *MAFA* is also expressed in non-photoreceptor retinal cells [19] as well as lens tissues [18], and can induce rods when ectopically expressed in mice [20].

We recently collaborated to compare lineage tracing events and prompt a new hypothesis that ancient mammals began to convert a large proportion of cone-fated progenitor cells to the rod cell fate [20]. This was proposed as part of the mammalian adaptation to the Nocturnal Bottleneck [20], a phase of evolution in the earliest proto-mammals where they avoided daytime predators by adapting to a nocturnal lifestyle [21, 22]. We speculated that changes in NRL expression or function may have been involved in capturing cone cells to the rod fate. A burst of evolutionary change in NRL peptide sequence, well conserved among mammals but less so outside the clade [20], suggests a change in NRL functions and/or roles in development, as does the differing capacity for Xenopus *Nrl* versus human *NRL* to induce lens fibre differentiation in the frog [16].

To facilitate the comparison of photoreceptor specification programs between dichromat (mammalian) and tetrachromat (early-branching vertebrate) models we challenged the hypothesis that the functional role of NRL has is conserved between mouse and zebrafish. To this end, we determined the outcome of *nrl* loss on a tetrachromat retina across various stages of ontogeny. We also tested the capacity for ectopic zebrafish or mouse *NRL* orthologs to override an established cone-specified phenotype in favor of a rod phenotype in transgenic animals expressing *nrl* in UV cones. Moreover, we developed lineage tracing tools to track these outcomes over time. Overall this paper identified both deeply conserved and diverging functional requirements for *NRL* between mice and zebrafish when considered over ontogeny.

## RESULTS

### Larval zebrafish require *nrl* to make rods early in ontogeny

Zebrafish *nrl* is the sole zebrafish ortholog to mammalian *NRL*; no paralogs have been identified [20]. The protein domains of zebrafish Nrl are conserved compared to mammals, though the zebrafish protein is longer in its primary sequence (Fig. 1B and S1). We used CRISPR/Cas9 targeted mutagenesis to create a loss-of-function allele of zebrafish *nrl*, and to assess its role in photoreceptor development (Fig. 1B and S1). We generated frameshift allele *nrl*^*ua5009*^ that was predicted to lack all major Nrl protein domains, and therefore was a putative null allele.

*nrl*^*ua5009*^ homozygous mutant zebrafish (referred to hereafter as *nrl*^*-/-*^) fail to produce rods at 4 days post fertilization (dpf) (Fig. 1C,D), contrasting wild type larvae that consistently produced a large abundance of rods by this time point. When examined using the rod-specific 4C12 antibody, we found that these *nrl* mutant zebrafish consistently contain zero rods within the entirety of the larval retina. *nrl*^*-/-*^ mutant zebrafish larval showed no overt abnormalities in retinal lamination (Fig. 1E,F).

Immunoblots on wildtype zebrafish detected Nrl protein at the expected size (predicted to be 44.2 kDa based on primary sequence). Zebrafish Nrl immunoreactivity presented as a doublet band (Fig. 1G) highly reminiscent of blots against mouse Nrl [23], where post-translational modification via SUMOylation has been determined experimentally to account for the doublet band [24]. SUMOylation of zebrafish Nrl is plausible, because the site of SUMOylation in human Nrl, residue K20 and surrounding residues [24], is exactly conserved in zebrafish (Supp Fig. S1A). Nrl protein was not detectable in *nrl*^*-/-*^ mutant retinas (Fig. 1G and see Supplemental Fig. S1 for full blots and quantification). Consistent with the lack of detectable protein, 5’RACE characterization showed no detectable alteration to splicing of the *nrl* transcript in *nrl*^*-/-*^ mutant larvae or adult retinas (Fig. 1H and see Supplement Fig. S1 for full blots). Thus 5’RACE discounts possible confounds to the prediction of a null allele in our mutants (e.g. that might occur via imagined splicing of cryptic exons).

In mice, lack of *Nrl* causes overproduction of an S-cone photoreceptors (in addition to loss of rod cells) [4]. To assess if one or multiple of the zebrafish cone subtypes are more abundant in *nrl* mutants, we determined the relative abundance of photoreceptors of wholemounted retinas. We found that rods were consistently absent in *nrl* mutants at 4 dpf (0 rods ±0 SEM, n = 37), whereas in wildtype larvae, within a 100×100µm region of interest (ROI), there were 141.45 rods ± 5.76 (SEM, n = 29) (Fig. 1D,I). We found that UV cones were significantly overproduced in mutants (mutant: 817.38 ± 26.60 (SEM); wildtype: 599.64 ± 21.45 (SEM); Mann-Whitney U test, p-value = 0.0001512, U stat = 112) (Fig. 1D,I). The excess UV cone abundance in *nrl* mutants was approximately equivalent to the normal abundance of rods in wildtype larvae, suggesting that cells otherwise fated to become rods might have become UV cones without *nrl*. Consistent with this, we did not detect a significant difference in blue cone or red cone abundance in *nrl* mutants (Fig. 1I)

We did not detect overt morphological or developmental consequences of *nrl* mutation in zebrafish beyond the photoreceptor phenotypes documented above, aside from a disruption in the normal development of the lens. By 2dpf, mutant zebrafish could be routinely distinguished from heterozygous or wildtype siblings by the presence of an occlusion in the lens (shown at 3dpf, Fig. S2). This is consistent with zebrafish *nrl* being robustly expressed in the developing lens fibre cells [14].

Controls for specificity of the above mutagenesis support that the frameshift lesion we induced in *nrl* is causal of the described photoreceptor phenotypes. Both phenotypes (reduction in rods, increase in UV cones) were recapitulated when *nrl* splicing was disrupted by morpholino (Supp Fig. S3). Furthermore, transgenic replacement of *nrl* was able to rescue the loss of rod cells on the *nrl*^*-/-*^ background (described immediately below and in Fig. 2C’’), arguing that our *nrl* mutant larvae possess all the molecular machinery required for producing rods (excepting *nrl* itself).

**Figure 2.**
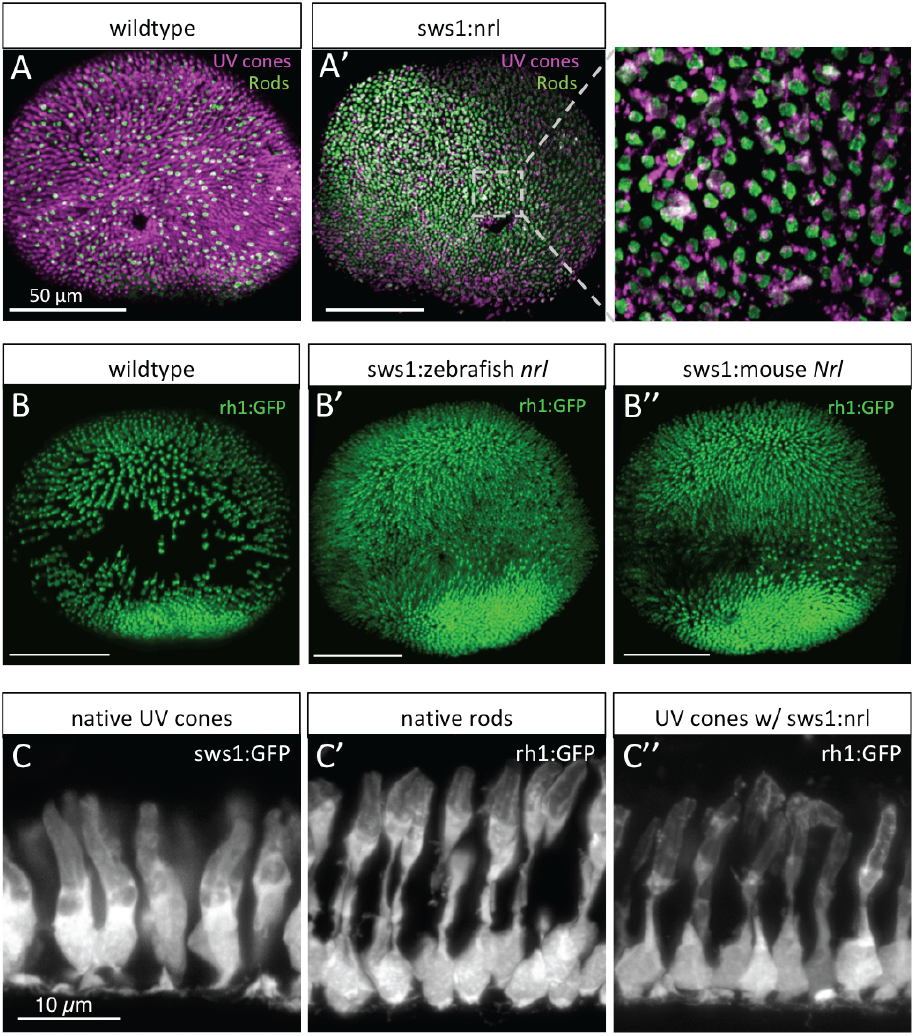
Zebrafish Nrl is conserved and sufficient to induce rod photoreceptors in zebrafish. **A**. Wildtype zebrafish larval retina in *en face* view has a dense forest of UV cone photoreceptor cells (magenta) and fewer rod cells (green) scattered throughout. **A’**. Ectopic expression of zebrafish Nrl in differentiating UV cones transmutes these cells to a rod cell fate as determined by 4C12 immunoreactivity (green; 4 days post-fertilization (dpf) larva). Inset: UV cones (magenta) are 4C12+ (green). **B**. Rods are more abundant in wildtype zebrafish by 6dpf. B” Expression of mouse Nrl reroutes cones to a rod cell fate (defined as GFP+ cells in *Tg[rh1:gfp]*) in a manner indistinguishable from zebrafish Nrl (B’). Note the increase in GFP+ rods apparent in both lines (B’ & B”) relative to wildtype retina. **C**. Cellular morphology of UV cones vs. rods (C vs. C’) are readily distinguishable by 7dpf in photoreceptor cells expressing GFP. **C’’**. Ectopic expression of zebrafish Nrl causes UV cones to take on a rod-like cell morphology. Larvae in C’’ are on a *nrl*^*-/-*^ null background (and thus lack native rod cells, described in Figure 3 to ensure the source of GFP+ rod-like cells visualized here is the UV cones ectopically expressing the transgenic Nrl.

### Zebrafish *nrl* is conserved and sufficient to convert UV cones to a rod-like fate

To test if zebrafish *nrl* is sufficient to induce a rod photoreceptor cell fate, we transgenically expressed *nrl* in developing zebrafish UV cones. We engineered transgenic zebrafish using regulatory sequences upstream of the *sws1* gene to drive expression of zebrafish *nrl*; this promoter has previously been characterized to drive expression exclusively in UV cones [25][20, 26-29], At 4 dpf, zebrafish with UV cones that expressed *nrl* showed 4C12 immunostaining (normally specific to rods) that colocalized with UV cones, as stained by anti-UV cone opsin (Fig. 2A). Thus, zebrafish *nrl* is sufficient to produce rod photoreceptors.

Ectopic expression of zebrafish Nrl in differentiating cones was sufficient to induce the rod opsin promoter and rod cell fate, such that *Tg[rh1:GFP]* larvae displayed a great density of GFP-positive cells (native rods and transmuted UV cones in Fig. 2B’) compared to wildtype siblings (Fig. 2B). Strikingly, the cellular morphology of the transmuted cones in *Tg[sws1:nrl]* retinas was nearly indistinguishable from native rods (Fig. 2C’’). Native rods can be readily distinguished from native cones (compare Fig. 2C’ *vs*. 2C) by their long thin morphology connecting apical outer segments to the basal cell body and nuclear compartment (Fig. 2C’), contrasting native UV cones that are more uniformly thick from their basal nucleus through their inner and outer segments (Fig. 2C). To confirm that the *rh1:GFP*-positive rod-shaped cells characterized in Figure 2C’’ were transmuted cones rather than native rods, these observations were made on an *nrl*^*-/-*^ background where native rods are absent (as described in Fig. 1; the data is reminiscent of Nr2e3 rescuing rod cells in *Nrl*^*-/-*^ mice [30]). Overall then, diverse markers (Fig. 2A,B,C) show that zebrafish Nrl is sufficient to induce rod characters in differentiating zebrafish photoreceptors.

To further challenge the hypothesis that zebrafish Nrl is functionally conserved with mouse NRL, we examined retinas from larval fish engineered to be similar to those above but expressing the mouse Nrl gene in developing cones. Mouse Nrl converted UV cones to rods in a manner indistinguishable from zebrafish nrl, revealed by the high density of rods in transgenic *Tg[sws1:Mmu*.*NRL-FLAG]* retina compared to wildtype siblings (compare Figure 2B’’ to B’ and their shared disparity compared to wildtype in panel B). Together with previous data from transgenic mice, where ectopic expression of Nrl homologs from mouse or chicken were sufficient to induce rod cell fate [20, 31], our data demonstrate that NRL homologs have a conserved capacity to induce rod cell fate across a diversity of vertebrates.

### Zebrafish cone cell lineage has high fidelity but can be induced to the rod cell fate

In adult transgenic zebrafish expressing zebrafish *nrl* in UV cones, UV cones were only detected near the CMZ (a region of retinal growth that continues to produce new retina into adulthood Fig. 3A, B). This could be due to the death of the UV cones shortly after they developed, or due to their conversion to a rod cell fate accompanied by the cessation of UV opsin production. To investigate this, we produced genetically encoded Cre-recombinase lineage tracing tools to assess cone cell fates.

**Figure 3.**
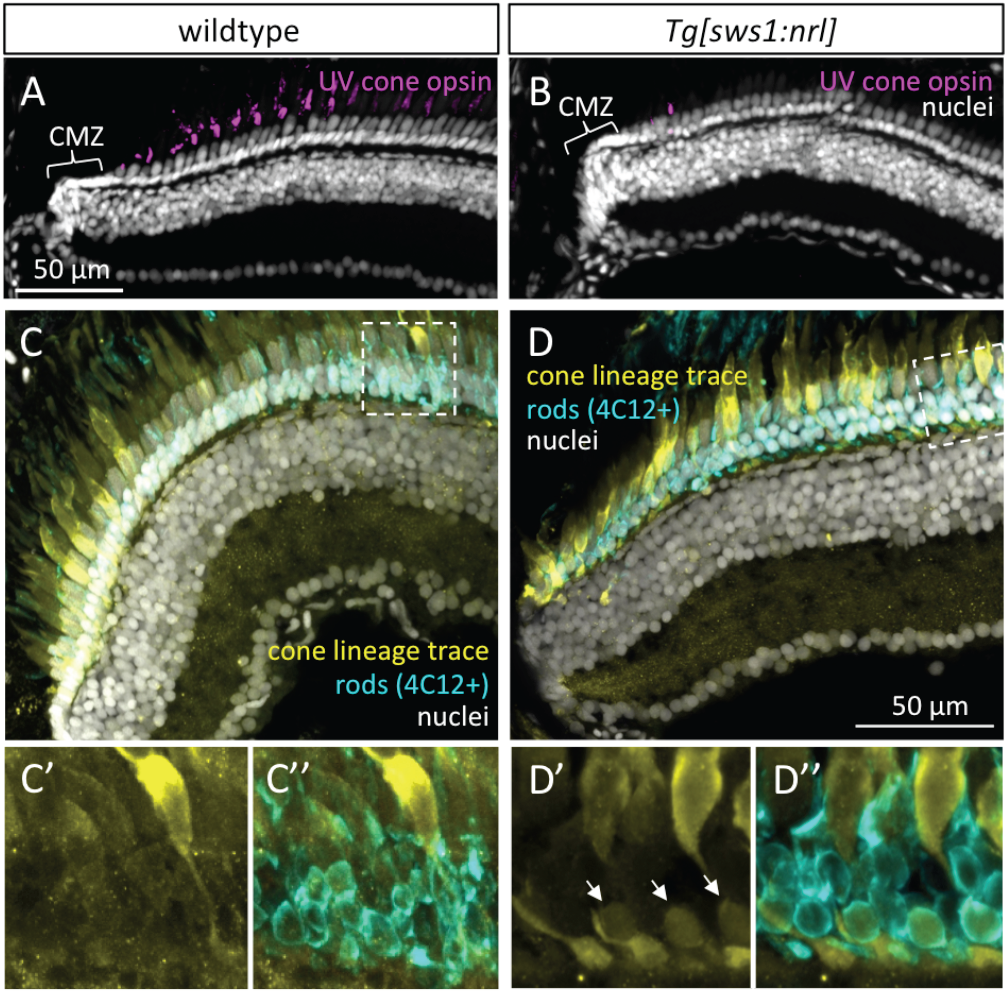
The cone cell lineage does not appreciably contribute to rod production in zebrafish but inducing ectopic Nrl shows it has this capacity. Adult zebrafish retina grows from proliferating cells in the ciliary marginal zone (CMZ), generating all cell types including regularly spaced UV cones (**A**). **B**. Following ectopic expression of Nrl in differentiating UV cones, adult retina is mostly devoid of UV cones, except sparse newly born UV-opsin+ cells near the CMZ. Panels A & B are anti-UV opsin immunohistochemistry (magenta) with nuclear counterstain. **C**. The adult zebrafish cone cell lineage gives rise to all cone cell types, and few other cell types are appreciably generated from that lineage, as detected by Cre-lox lineage tracing via a cone-transducin-α (*gnat2*) Cre driver line. No history of *gnat2* expression (yellow) is detectable in rod cells (4C12+), despite the lineage trace reporter being abundant in all other photoreceptors (cones). C’ & C” are alternate views of dashed box outlined in panel C. See also associated Supplemental Fig. S4. **D**. The fate of UV-opsin+ cells, that are absent from mature retina in (B), includes their transmutation into rods. Note a subset of rod cells (arrows, identified as 4C12+ and with nuclei in the basal-most layer of the ONL) show a history of *gnat2* expression (yellow), indicating they were generated from the cone lineage.

To follow the fate of UV cones that ectopically express nrl into adulthood, we used a paradigm of genetically encoded lineage tracing wherein <u>all cells emanating from the cone lineage will permanently express fluorescent reporter regardless of their subsequent cell fate</u> (Fig. S4A,C). We drove transgenic expression of Cre recombinase under control of regulatory sequences for *gnat2* (cone transducin α) to induce Cre expression in all subtypes of developing cones [29, 32]. We bred this transgenic (Tg) line to two lox-mediated reporter lines, Zebrabow Pan, Freundlich [33] (Fig. S4A) and ubi:Switch [34, 35] (Fig. S4C). In this cone lineage tracing line, zebrafish possessing wildtype *nrl* had no rods that were observed to originate from the cone cell lineage (i.e. none expressed *gnat2*:Cre-mediated fluorescent reporter; 8/8 fish with Ubi:Switch, 1/2 fish with Zebrabow, Fig. 3C & S4E).

The fate of disappearing UV cones (the UV cones expressing ectopic Nrl in our Tg fish) was examined by incorporating these same cone lineage tracing reporter constructs into *Tg[sws1:nrl]* fish. Contrasting the results from wildtype zebrafish (Fig. 3C), cone lineage tracing demonstrated that *Tg[sws1:nrl]* fish possessed rods expressing the cone lineage reporter (8/8 fish with Zebrabow) (Fig. 3D & S4F, arrowheads). The characterization of cells from the cone lineage as being rods, in *Tg[sws1:nrl]* fish, was based on both the basal location of their nuclei within the ONL and their immuno-colocalization with the rod marker 4C12 (Fig. 3D).

In 1 of 2 lineage trace zebrafish with wildtype *nrl*, using the Zebrabow paradigm, we noted a total of 27 lineage-trace-reporter-positive rods, in 5 clumps, in the oldest parts of the retina near the optic nerve head (Fig. S4E); in the same fish, there was also expression of lineage reporter in other cell layers, including in some bipolar cells and ganglion cells. Considering the rarity of these rod cells and their presence in only a single fish (and only from one of two reporter lines), we suggest they are attributable to stochastic Cre-like DNA recombination events or spurious expression of the transgenic construct. The robust and abundant expression of cone lineage tracing reporter in rods only occurred in animals that also expressed Nrl in UV cones, suggesting that expression of Nrl in zebrafish UV cones is sufficient to reprogram them to a rod phenotype. These conclusions were supported in larval fish (6/6 retinas examined; Supp Fig. S4D) using the Zebrabow reporter line. This confirms that rod development in zebrafish proceeds in a straightforward manner that does not incorporate the cone lineage, i.e. as we had assumed-it-to-be prior to learning that mice utilize a more circuitous route and also produce rods via the cone progenitor lineage [20].

### >Nrl is dispensable for the specification of rods in adult zebrafish

To assess the impact of *nrl* loss on adult photoreceptors, we examined the photoreceptor composition of adult *nrl*^*-/-*^ mutant fish. Surprisingly, zebrafish adults with homozygous *nrl* mutation produce abundant rod photoreceptors (Fig. 4B-C).

**Figure 4.**
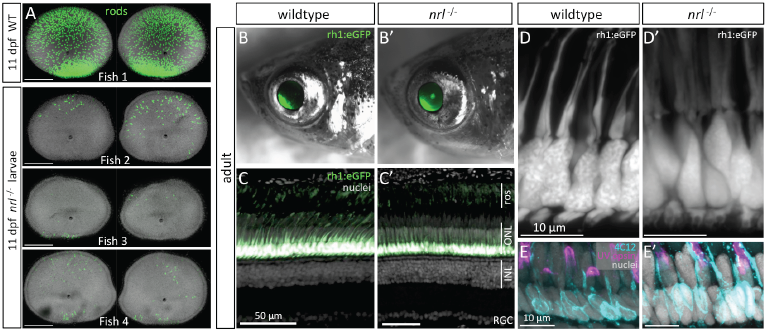
Nrl is dispensable for rod specification in adult zebrafish. **A**. Monitoring for appearance of GFP+ rods during ontogeny of *nrl*^-/-^ larvae directed our attention to 11 days post-fertilization (dpf), where rods are sporadically detectable, but varied between individuals and between eyes of the same individual. Wildtype (WT) eyes at top with abundant rods provided for context. **B-D**. Adult *nrl*^-/-^ zebrafish possess a large abundance of rods indistinguishable from WT, such that GFP+ rods are obvious in intact animals (B) or retinal cryosetions (C). Adult *nrl*^-/-^ retina showed normal distribution of rod outer segments (ros) apical of rod cell bodies in the outer nuclear layer (ONL, also in panel D). Inner nuclear layer (INL) and Retinal Ganglion Cell layer (RGC) are overtly normal (quantified in Fig. S5G,H). **E**. Immunolabelling with rod-specific 4C12 and anti-UV-opsin confirms presence of rods and normal UV cones, respectively, in adult *nrl*^-/-^ retina.

To affirm that the presence of rods in adult *nrl*^*-/-*^ fish represented a difference based on ontogenetic stage (rather than a stochastic difference between individuals), and to characterize when rods first appear in the *nrl*^*-/-*^ retina, we sought to assess individuals through their development. Breeding *Tg[rh1:eGFP]* into the *nrl*^*-/-*^ background allowed us to monitor for the appearance of GFP-positive rod cells in the eye of developing larvae, and directed us to focus our characterization on 11 dpf. The benchmark comparator is wildtype larvae at 11dpf, where retinas consistently had a large abundance of rods (thousands of rod cells per eye), including a concentration of rods in the ventral region and a large density of rods throughout all other retina regions (e.g. Fish 1 at the top of Fig. 4A). Retinas from *nrl*^*-/-*^ larvae at 11 dpf possessed a scattering of rods (∼10-100 rod cells per retina, Fig. 4A), contrasting younger *nrl*^*-/-*^ larvae where rods were never observed. Notably, the abundance and location of the rods varied considerably between individuals, and between the two eyes within individuals (three representative examples displayed in Fig. 4A, and such variation was apparent following examination of dozens of larvae). The stochastic nature of rod distribution in larval *nrl*^*-/-*^ fish is no longer apparent in adults, where the large abundance of rods is not different from wildtype animals (Fig. 4C and quantified in Supp Fig. S5G,H). We estimate that rods can be produced using a mechanism independent of *nrl* beginning late in the ontogeny of larval zebrafish, stochastically appearing at about 11 dpf.

In adult retinas, all the rod photoreceptors markers that we tested each affirmed the presence of normal rods in *nrl*^*-/-*^ null zebrafish. Rod opsin transcript (*rh1*) is appropriately localized and abundant in adult *nrl*^*-/-*^ retina as determined by *in situ* hybridization on cryosections and by RT-qPCR (Fig. 5). Rods in adult *nrl*^*-/-*^ retina were also immunopositive for antibody 4C12 (Fig. 4E’) and robustly express the rod-specific transgene *Tg[rh1:eGFP]* (Fig. 4B,C). The morphology of the *nrl*^*-/-*^ rod cell bodies is normal at the level of confocal microscopy (Fig. 4D), and the rod outer segment is normal at the level of ultrastructure (Fig. 6A) showing freely-floating disks indistinguishable from wildtype.

**Figure 5.**
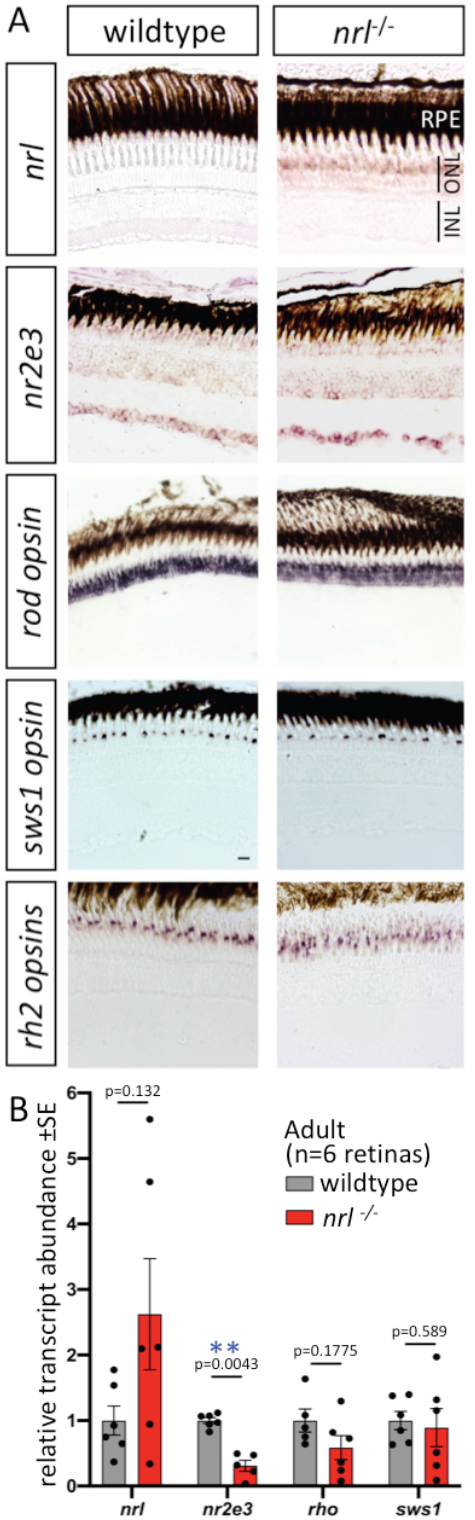
Adult retina of zebrafish *nrl*^-/-^ mutants show changes in abundance of Nrl target gene *nr2e3* and in *nrl* itself. Relates to Figure 1 regarding confirmation that Nrl is disrupted, and Figure 4 regarding lack of expected photoreceptor phenotypes, in adult retina. **A**. Gene expression determined by *in situ* hybridization on cryosections of adult zebrafish retina. Expression of *nrl* is highly enriched in the outer nuclear layer, consistent with the site of phenotypes when it is disrupted. Abundance of *nrl* transcript is higher in *nrl*^-/-^ frameshift mutant retina (confirmed in panel B), suggesting an auto-regulatory negative feedback loop controlling it own abundance. Alterations to the abundance of *nr2e3*, a downstream target of Nrl, are equivocal when measured by *in situ* hybridization. The levels and distributions of transcripts encoding rod (*rh1*) and cone (*sws1* and *rh2*) opsins were not detectably different in *nrl*^-/-^ mutant retina compared to wildtype. Scale bars are 10 μm. **B**. Transcript abundance in adult neural retina determined by RT-qPCR confirms an increase in *nrl* abundance in zebrafish bearing a frameshift null allele in *nrl*. A downstream transcriptional target of Nrl, *nr2e3*, was 70% reduced in abundance in *nrl*^-/-^ mutant retina compared to wildtype (p<0.01; n=5-6 individuals per genotype). Opsin abundances in adult retinas were not markedly different between genotypes.

**Figure 6.**
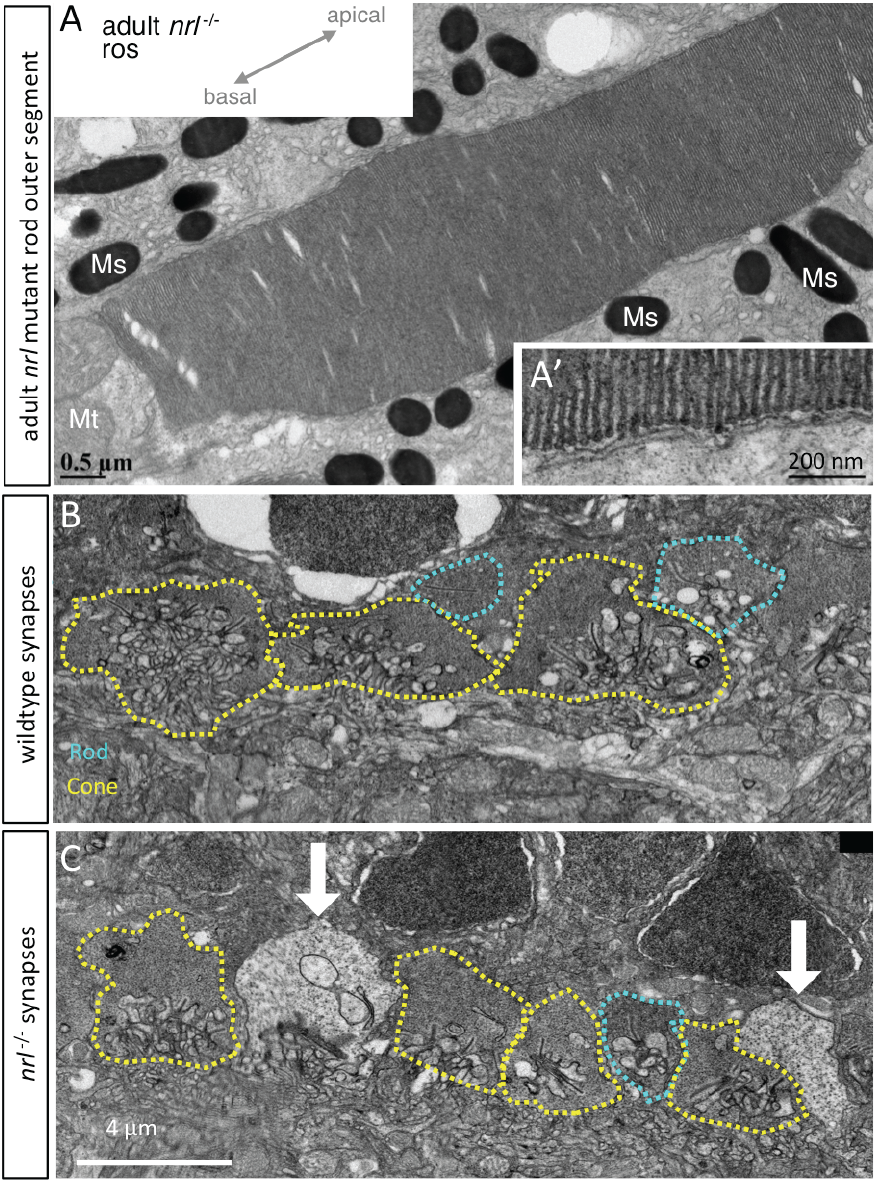
**Rod outersegments of *nrl*^-/-^ adult zebrafish appear normal,** demonstrating the expected hairpin end of floating disks within the outer segment that are non-contiguous with the outer cell membrane **(A’)**, diagnostic of rod cell identity. **B**. In wildtype adult zebrafish photoreceptor synaptic terminals include rod spherules (teal dotted line) that are morphologically distinguishable from cone pedicles (yellow). **C**. In *nrl*^-/-^ adult retina the cone pedicles appear normal, and rod spherules appear normal but are very sparse; instead electron-lucent terminals (white arrows) uniquely appear and may represent an Nrl-dependent defect in rod synapse maintenance. Various characters of photoreceptor terminals are quantified in Fig. S6. Mt, mitochondria; Ms, melanosomes.

An important aspect of concluding that Nrl is dispensable for rods in adult zebrafish is assessing whether any Nrl function is retained in adult *nrl*^*-/-*^ fish. A downstream effector of Nrl, *nr2e3*, was found to be ∼70% less abundant in adult *nrl*^*-/-*^ retina (p=0.0043; Fig. 5B), consistent with Nrl immunoreactivity being abrogated in immunoblots from adult mutant retina (Fig. 1G, Supp Fig. S1D), and supporting that this is a null allele. Characterizing *nrl* transcripts in *nrl*^*-/-*^ retina further supported a potent disruption, insomuch that *nrl* transcript abundance was increased ∼2.5 fold in adult *nrl*^*-/-*^ retina (not significant; Fig. 5B); we infer that *nrl* may negatively regulate its own abundance (either directly or indirectly) and the absence of functional Nrl protein removes this negative autoregulation. The increased abundance of *nrl* transcript was also apparent via *in situ* hybridization on retina sections, and offers good support that *nrl* expression is highly enriched in the outer nuclear layer (Fig. 5A), and exactly consistent with the cellular location of the *nrl*^*-/-*^ phenotype (ONL containing the photoreceptor nuclei). The mutations in *nrl* also cause the abrogation of rod cells in larval fish and this exactly matches phenotypes predicted from *Nrl* knockout mice. Overall, disruption of downstream targets and loss of Nrl immunoreactivity in adult retina strongly match the prediction that our engineered frameshift mutation leads to a *nrl*^*-/-*^ null allele. Therefore, zebrafish are able to produce rod photoreceptor cells in their adult stages via a pathway independent of *nrl*.

### Lack of obvious phenotypes in cones or other retinal cells of adult nrl^-/-^ zebrafish

In light of the cone photoreceptor phenotypes in *Nrl* knockout mice, including an increase in sws1 cones (“S-cones”) [4] that we had also observed in larval *nrl*^*-/-*^ zebrafish (Fig. 1), we characterized cone photoreceptors in adult *nrl*^*-/-*^ zebrafish. The highly organized cone mosaic in adult zebrafish retina, wherein cone spectral subtypes are positioned into repeating rows with high precision and fidelity [36-40], ensures that any notable disruption to cone photoreceptors is obvious. No abnormal cone phenotypes were apparent in adult *nrl*^*-/-*^ retina. In particular, an overtly normal cone mosaic was apparent in adult *nrl*^*-/-*^ retina based on position of the nuclei (Fig. S5A-F), with zpr1+ double cones flanking a single cone nucleus to form repeating pentameres of cone nuclei across the retina (Fig. S5A,B). Lws cones and sws2 cones (red- and blue-sensitive cones) were spaced evenly across the adult *nrl*^*-/-*^ retina in their expected patterns, as detected by anti-lws-opsin and *Tg[sws2:mCherry]*, respectively (Fig. S5C-F). Green-sensitive cones were distributed evenly across the adult *nrl*^*-/-*^ retina as determined by *in situ* detection of *rh2* opsin (Fig. 5A). Immunolabelling and *in situ* detection of *sws1* opsin demonstrates adult *nrl*^*-/-*^ retina possess a normal spacing of UV cones (Fig. 4E & 5A, respectively). Tracing the cone lineage in adult *nrl*^*-/-*^ retina suggested a normal generation of cone subtypes (e.g. with no rod cells from the cone lineage) with the expected diversity of elaborated cone morphologies (Fig. S5I).

Other retinal phenotypes associated with *Nrl* knockout mice, including rosettes or other defects in retinal lamination [5, 41], were not observed in retinas of *nrl*^*-/-*^ zebrafish (Fig. 4C). Regardless, we explored for more subtle disruptions by quantifying cell abundances in radial retinal sections. In neither central retina nor peripheral retina is any difference in cell abundance detectable including for rods, cones, retinal ganglion cells, horizontal cells or other inner nuclear layer cells (n=6 fish per genotype; Supp Fig. S5G &H).

### Requirements for Nrl in maintaining the adult zebrafish rod synapse

We noted that ultrastructural aspects of photoreceptor synaptic terminals were abnormal in adult *nrl*^-/-^ zebrafish. Cone pedicles were recognizable in wildtype and *nrl*^-/-^ retina due to their larger size, large abundance of synaptic ribbons and somewhat electron-lucent appearance relative to wildtype rod spherules (Fig. 6B). Ribbon length, vesicle density and pedicle area in cones showed no dramatic differences between genotypes (Supp Fig. S6). However normal rod spherules were rarely observable in adult *nrl*^-/-^ retina (Fig. 6C, S6B), although when present they appeared normal in the aforementioned characters (Supp Fig. S6C-E). Unique to adult *nrl*^-/-^ retina, we noted a population of photoreceptor terminals that were broadly cone-like but extremely electron-lucent such that we designated them as ‘white synapses’ (Fig. 6C, found abundantly in 4/5 *nrl*^-/-^ fish examined). Considering the discrepant large abundance of rod cells vs. the paucity of normal rod spherules in adult *nrl*^-/-^ retina, we infer these white synapses likely belong to *nrl*^-/-^ rod cells. Indeed confocal characterization of GFP-filled rod spherules (e.g. visible in bottom of Fig. 4D) showed that the synaptic invagination was larger (perhaps more ‘cone-like’) in *nrl*^-/-^ rod spherules compared to sibling rod spherules (Supp Fig. S6F). Compared to wildtype rod spherules, *nrl*^-/-^ white synapses had ultrastructure that was somewhat more cone-like (larger size, longer ribbons, quantified in Supp Fig. S6C-E).

We found that the cone nuclei of wildtype and *nrl*^*-/-*^ adults were similar in chromatin texture and electrolucency (Fig. S7C,D). However we found that mutant rod nuclei had a mottled, though frequently still electron-dense appearance (Fig. S7A,B). In the mouse, wildtype rods have a characteristic electron-dense arrangement of heterochromatin in the centers of their nuclei [5, 42], whereas cone nuclei have a mixed arrangement of hetero- and euchromatin, leading to a mottled appearance. These two nuclear phenotypes are conserved in the zebrafish [43]. Thus, *nrl*^*-/-*^ zebrafish rods may be thought of as exhibiting a cone-like chromatin appearance.

In sum, rod photoreceptors in adult *nrl*^-/-^ zebrafish are inferred to have electron-lucent and somewhat cone-like synaptic terminals, perhaps suggesting they have defects in synapse maintenance or differentiation. Rod photoreceptors in adult *nrl*^-/-^ zebrafish also have somewhat cone-like nuclear condensation at the ultrastructural level. However rod photoreceptors in adult *nrl*^-/-^ zebrafish are specified as rod cells such that the diagnostic features of rod outer segments (rod opsin expression and disk ultrastructure) appear normal.

## DISCUSSION

Our data describe a conserved role for Nrl early in zebrafish ontogeny that recapitulates its well-researched master-regulatory role in mice, with Nrl being both necessary and sufficient for rod photoreceptor cell development. However, our data surprise by revealing that Nrl is dispensable for rod cell specification in adult zebrafish.

The rods present in adult *nrl* mutants have gene expression and ultrastructural differences from normal rods, including cone-like nuclei and synapses, indicating that *nrl* is involved in, but not solely responsible for, rod development in adult zebrafish. We also evaluated the capacity for zebrafish *nrl* to over-ride the cone specification program in maturing UV cones, and found that the erstwhile UV cones eventually adopted an unambiguous rod-like phenotype, indicating a conserved capacity for zebrafish *nrl* to induce rod characters. Indeed this capacity to induce rod characters was indistinguishable in transgenic fish expressing mouse or zebrafish homologs of *Nrl*.

Appreciating the mechanisms whereby the transcription factor NRL specifies rod *vs*. cone photoreceptor cells is foundational knowledge to at least five disparate fields of evolutionary and biomedical research. First, this work will be of interest to theorists seeking to understand the mechanisms of how novel cell types arise over evolutionary time. Proposals for these mechanisms have benefited from a focus on photoreceptors, which have rapidly diversified early in vertebrate evolution, and the role of Nrl in mouse photoreceptor specification has been an instructive case study in cell type evolution [44]. The existence of Nrl-independent phenotypes in zebrafish rods may indicate an additional cell type diversification among rods which can be further mined for insight. Second, mutations in the *NRL* gene (along with its downstream effector *NR2E3*) are causal of blinding disorders that remain unchecked [45, 46]. Third, *Nrl*^*-/-*^ knockout mice have broadly imposed themselves upon animal modelling of Ophthalmology, often being used as a (relatively artificial) proxy for the cone-rich macular region of human retina. Fourth, gene therapy strategies that disrupt *NRL* show substantial promise in mouse models as a cure for retinitis pigmentosa and other rod degenerative disease – a mechanism to save degenerating rods by converting then into a cone-like state [47]. Finally, stem cell therapies to repair vision loss have demonstrated great potential but must now overcome the hurdle of regenerating cones (rather than rods) to repair daytime and high-acuity vision, and *NRL* is at the heart of the gene network driving this cell fate switch. Appreciating NRL function beyond nocturnal mice begins to fill a substantial and influential knowledge gap: It is surprising that the function of Nrl had remained untested in zebrafish, despite zebrafish emerging as the premier genetic model of vertebrate photoreceptor development and regeneration.

Moreover, we and collaborators recently proposed that evolution in Nrl’s function and/or regulation are prime candidates for a proximate mechanism in the evolutionary success of early mammals as they adapted to survive the nocturnal bottleneck [20]. Comparative lineage tracing between mice and zebrafish demonstrated that a majority of rod cells arise from an unexpected source in mice – the cone progenitors – whereas cone lineages only gave rise to cones in zebrafish. The lineage tracing results in mice were bolstered by analyses of transcripts and protein in developing rods that revealed vestiges of sws1 cones [20]. That taxonomic comparison suggested a developmental innovation had contributed to evolving the rod-rich retina of mammals, and we proposed changes at the Nrl locus of early mammals as a potential driver of this adaptation. The proposal aligns well with previous data showing ectopic expression of Nrl in mouse retinal progenitors as being sufficient to drive the rod cell fate [31].

### Lineage tracing of adult rod photoreceptors using *gnat2*-induced reporter in the absence of ectopic *nrl* expression

We previously demonstrated that larval zebrafish rods do not have a history of *sws1* expression [20], using a Gal4/UAS-derived technology. Zebrafish silence the UAS promoter as they age and over generations, precluding the use of that arrangement of genetically encoded lineage tracing constructs for this study. Here, we found that another cone gene, *gnat2*, also did not report expression in any rod in zebrafish larvae using a Cre/Lox lineage tracing system (Fig. S4D), consistent with and extending the previous study concluding that larval zebrafish rods do not exhibit a history of expressing cone genes. While the *gnat2:cre* lineage tracer robustly reported rods in conjunction with ectopic *nrl* expression in UV cones (Fig. 3 & S4), we noted 5 clusters of lineage-traced rods, along the length of a retinal section from CMZ to optic nerve head, in one animal without transgene-induced ectopic *nrl* expression (Fig. S4E, arrowheads). In animals of both genotypes, we noted occasional labeling of other cell types (e.g., Fig. S4F, a bipolar cell). We consider it likely that the 5 clusters of lineage traced rods represent 5 clonal populations of rods with one-time spurious expression of *gnat2:cre* during their development.

### Zebrafish *nrl*^*-/-*^ rods, and cone-rod transmutations in nature

Close examination by electron microscopy of the adult *nrl*^*-/-*^ rods suggested that the mutants do not make typical rod synapses; across two mutant animals, we found a single synapse that was clearly rod-like: electron dense relative to neighbouring cone photoreceptors, a single synaptic ribbon that was longer than nearby cone ribbons, and placed either between cone synapses, or positioned slightly scleral within the synaptic layer, consistent with previous zebrafish synapse characterization [43]. The remaining synapses appeared to be either cone synapses, or in one mutant animal, “white” synapses. The “white” synapses were positioned like cones within the synaptic layer, interspersed among cone synapses. There were not enough of them to fully account for the lack of obvious rod synapses; if they belonged to the rods, then other synapses, perhaps more overtly cone-like synapses, likely did as well. This is reminiscent of the synapses of lamprey photoreceptors. At least one species of Lamprey have cells with the physiological characteristics of rods: the ability to respond reliably to single photons of light, sluggish responses to stimulation relative to the more cone-like lamprey photoreceptors, and the ability to send their signals to the cone-like photoreceptors [1, 3]. However, lamprey rod-like cells also have cone-like characteristics; *Mordacia mordax*, a nocturnal lamprey with a single photoreceptor with rod-like physiology, has plasmalemma invaginations in the outer segment that mean it does not have rod-like free-floating membrane discs, but instead has a cone-like morphology [2, 48]. *Petromyzon marinus* has rod-like synapses associated with the physiologically rod-like cell, but these can have up to 4 synaptic ribbons, and the ribbons do not appear to differ in length between photoreceptor types [49]. Outside lamprey, the teleost deep-sea pearlside *Maurolicus muelleri* has photoreceptors deemed “true” rods (rhodopsin-expressing; minority) and rod-like cones (green opsin-expressing; majority), and the synapses of the rod-like cones were smaller (rod-like), but had multiple synaptic ribbons per terminus (cone-like) [50]. It was proposed that the rod-like cones of *M. muelleri* are “transmuted” cones, in the sense of Walls [22].

The mottled chromatin texture of the *nrl*^*-/-*^ mutant rod (Fig. S7) is consistent with the mottled chromatin of the cone-like photoreceptors produced in the *NRL*^-/-^ mouse [5], and reminiscent of the texture of zebrafish cones (Fig. S7, and [43]). The dense heterochromatin of wildtype rods is possibly a solution to improve photon transmission and thus increase sensitivity of rods in dim light conditions, and is particularly consistent among nocturnal animals [51]. We note the similarity of the *nrl*^*-/-*^ zebrafish rod nuclei, which appear to be rods despite ultrastructural cone-like similarities, to the nuclei of the rod-like photoreceptors in *M. mordax* (see Fig. 7A, B of Collin and Pottert, 2000 [48]) and the nuclei of the rod-like photoreceptor of *P. marinus* are clearly mottled and nearly indistinguishable from the cone-like photoreceptor nuclei (see Fig. 12, 13 of Dickson and Graves, 1979 [49]. In the avian lineage, which unambiguously has rod-like photoreceptors but which has also lost NRL, electron microscopy of the common buzzard (*Buteo buteo*) suggested that all photoreceptors have mottled chromatin, although this was not explored in depth [52]. In the deep-sea pearlside *M. muelleri*, the nuclei of both the true rods and rod-like cones had obvious mottled chromatin [50]. It was not reported whether *M. muelleri nrl* was present in the transcriptome data. Furthermore, it is possible that examination of photoreceptor synapses and nuclei could be used to probe for cryptic or suspected transmutation events.

Thus, there are numerous examples of animals bearing rods with some cone-like aspects, which align with scenarios where an originally cone-like cell may have become rod-like. This would imply that the *nrl*^*-/-*^ zebrafish rods use cone machinery, or are derived incompletely from cells that started as cones. Future work might assess whether the *nrl*^*-/-*^ zebrafish rods deploy cone phototransduction machinery, as cone-rod transmutation events seem to leave various species using a mixture of cone and rod proteins for phototransduction. Our initial examination of this, with the gnat2:cre lineage tracing construct, did not label zebrafish *nrl*^*-/-*^ rods, suggesting they do not derive from post-mitotic cone-fated cells, and furthermore suggesting that these rods do not employ gnat2 (a cone phototranduction gene); however many other cone phototransduction genes ought to be assessed to determine whether there is a blended phenotype between cones and rods.

## Conclusion

The classic interpretation is that NRL is the absolute master regulator of the rod photoreceptor cell fate; we demonstrate here that this role is deeply conserved, and yet not completely conserved, in an intriguing manner outside of mammals. Despite the canonical requirement for NRL being apparent in larval zebrafish, adult zebrafish lacking Nrl were shown to specify and produce an abundance of rod photoreceptors. The *nrl*^*-/-*^ rods show only subtle ultrastructural and transcriptional differences from wildtype rods. The unexpected tolerance of adult zebrafish rods for the absence of nrl suggests that larval, not adult, zebrafish rods are best suited for biomedical modelling of mammalian rod dystrophies.

Across ontogeny, nrl expression is sufficient to induce a robust rod phenotype in developing cone photoreceptors, suggesting that cone photoreceptors remain competent to transmute to rods throughout the life of the animal. This has implications for the mechanisms enabling cone-to-rod transmutations found in various vertebrate lineages, including in early mammals.

Zebrafish retain ancestral vertebrate retina traits that have been lost in mammals, including the original complement of 4 cone subtypes and 1 rod. The unexpected difference between fish and mouse in nrl requirements reinforces the need to explore the developmental genetics of retinal cells across a range of vertebrates if we are to build a comprehensive understanding of retinal development and evolution.

## Acknowledgements

We appreciate technical assistance of Daniela Roth and Evelyn Free. Transgenic fish were generously shared by Marc Ekker (*Tg[ubi:switch]*). James Fadool kindly shared antibody 4C12, and Rachel Wong shared Gateway plasmid containing *gnat2* promoter region. We are grateful for advice and discussions with David Eisenstat, Paul Melancon, Joe Corbo and Sally Leys, and for comments on an earlier version of the manuscript by Anand Swaroop. The Natural Sciences and Engineering Research Council of Canada (NSERC) funded studentships to APO, EMD and KC and operating grants to WTA.

## Author Contributions

Conceptualization, A.P.O & W.T.A; Investigation, A.P.O, G.J.N., E.M.D., S.D.B., K.C.; Writing – Original Draft, A.P.O., W.T.A; Writing – Review & Editing, A.P.O, G.J.N., E.M.D., S.D.B., K.C., W.T.A.; Supervision, W.T.A. Project Administration, W.T.A; Funding Acquisition, W.T.A.

## MATERIALS AND METHODS

### Animal Ethics

All protocols involving zebrafish husbandry or experimentation were approved by institutional ethics Committees at the University of Alberta (protocol # AUP00000077), as overseen by the Canadian Council on Animal Care.

### Transgene Construct Cloning

Transgene expression plasmids were created using by incorporating gBlock Gene Fragments (from IDT, Integrated DNA Technologies, Coralville, Iowa) into the Multisite Gateway Cloning system. Zebrafish and mouse *nrl*, (appended with attB1 and attB2r flanking sequences, including a Kozak sequence, and a N-terminal 3xFLAG tag on Mouse Nrl), were both ordered as a gBlock Gene Fragment. Where applicable, silent mutations were engineered into the CDS of the various peptides in order to optimize for zebrafish codon bias and to circumvent nucleotide composition/ complexity requirements; in all cases, the predicted peptide produced is wholly wildtype aside from the relevant N-terminal tag. Received gBlocks were then processed for Multisite Gateway Cloning using the zebrafish Tol2Kit reagents [53] and Gateway system (Life Technologies), and were ultimately recombined with regulatory sequences of p5E-sws1 [26] and a p3E-polyA sequence into an expression vector. The expression vector backbone pDestTol2CG2 has a cmlc:eGFP reporter that drives eGFP expression in the heart muscle cells, to aid in identifying transgenic fish. Transgene expression constructs driving Cre recombinase expression were similarly assembled and cloned as above in front of p5E-gnat2 promoter [29, 32]; The p5E-gnat2 gateway plasmid was a kind gift from Dr. Rachel Wong of the University of Washington, USA.

### Morpholino and Plasmid Injections

Morpholino was injected into single cell embryos bearing rh1:eGFP and either sws1:nfsb-mCherry or sws2:nfsb-mCherry were injected with 10ng of standard control (CCT CTT ACC TCA GTT ACA ATT TAT A) or *nrl* splice-blocking morpholino (ACG TGT CAG ATC ATA CCT GTG AAG T) (Genetools, LLC, Philomath, Oregon), delivered to the yolk as a 5nL bolus. Morpholinos were first suspended in water and then diluted into 0.1M potassium chloride with 0.1% phenol red added to assess injection success. Morphants were reared to 4dpf and then processed for retinal mounting and imaged.

Transgene construct injection mixtures were prepared by diluting 750ng plasmid construct and 250ng Tol2 mRNA into 0.1M potassium chloride with 0.1% phenol red to a total volume of 10 µL. These solutions (5-10nL) were delivered to the yolk of the single cell embryo. Injected animals were raised to 2 dpf, and then inspected for eGFP-expressing heart cells, the marker of pDestTol2CG2 plasmid presence. Larvae with GFP-expressing heart cells were reared to adulthood, and germline transmission of transgenes identified in the subsequent generation.

### CRISPR Mutagenesis

To engineer allele *nrl*^*ua5014*^, a mixture of three guide RNAs (gRNA) targeting the first coding exon of zebrafish nrl were delivered. These gRNA were designed in Geneious R9.1.7 and synthesized per Gagnon and colleagues [54], but using mMessageMachine SP6 kit (Invitrogen). The gRNAs were mixed with protein Cas9 (NEB), allowed 5 minutes at 37°C to assemble into ribonucleoprotein complexes, and the mixture was microinjected into the cell of zebrafish embryos at the 1 cell-stage. Animals at 2dpf were sacrificed for high-resolution meltcurve (HRM) analysis of *nrl* cutting (STAR methods for primers; methods as previously described [55, 56].

For *nrl*^*ua5009*^, the 5’-most guide RNA used above, and a second guide RNA targeting eGFP (CR.GFP.GA5’.break, STAR methods) were mixed together and allowed to complex with protein Cas9. This mixture was microinjected into the cell of 1 cell-stage zebrafish embryos, bearing the ubi:eGFP transgene (STAR Methods), which promotes eGFP expression in all cells. At 2dpf, a subset of animals with disrupted eGFP were checked for nrl mutation, and the remainder of eGFP-disrupted fish were reared to adulthood and visually screened for germline transmission of mutant eGFP (carriers had no eGFP fluorescence, but the transgene bears a cardiac-expressed RFP marker cassette [26]). Fish with germline CRISPR editing were then checked for germline-transmitted nrl mutation, the *nrl*^*ua5009*^ allele was recovered, and ubi:eGFP^mutant^ was bred out in subsequent generations.

### Genotyping RFLP

Genotyping the *nrl*^*ua5009*^ allele leveraged a *TaqI* restriction enzyme cut site unique to the mutant allele. PCR amplification of *nrl*, using primers listed in table, was performed from genomic DNA. PCR parameters: 96°C 2min, 35x [56°C 15s, 72°C 1min, 96°C 15s], 72°C 5 minutes. PCR products were digested with *TaqI* restriction enzyme and analyzed by gel electrophoresis; the ∼675bp amplicon from the *nrl*^*ua5009*^ allele is cloven into two products. The *nrl*^*ua5014*^ allele was genotyped using these same PCR conditions and then resolving its larger amplicon, caused by the 109bp CRISPR-induced insert, via gel electrophoresis.

### 5’RACE to characterize mutant *nrl* transcript

RNA was purified from adult zebrafish retina or pools of 4dpf larvae using an RNeasy Minikit (Qiagen catalogue # 74104. Hilden, Germany). RNA sample quality was determined by Bioanalyzer 6000 RNA nano chip (Agilent catalogue # 5067-1512. Santa Clara, CA, USA), selecting for samples with RIN numbers >9. 5’ RACE was performed using a Clontech SMARTer RACE 5’/3’ kit (Takarabio catalogue # 634858. Kusatsu, Shiga Prefecture, Japan) with a primer in the third coding exon of *nrl* transcript ENSDART00000168271.3 (GATTACGCCAAGCTTTCTGTTCAGCTCGCGCACAGACAGACTC). Products were characterized on a 1% agarose gel and then purified using a Nucleospin Gel and PCR cleanup kit (Machery-Nagel # 740609.10. Düren, Germany). In fusion Cloning was performed using the Clontech SMARTer RACE 5’/3’ kit and identity of the 5’RACE products were verified, including appropriate presence/absence of the engineered mutation in allele ua5009, by sequencing using standard M13 primers.

### RNA Isolation and Quantitative real time polymerase chain reaction (qRT-PCR)

Neural retinae were dissected from adult zebrafish dark adapted for 12 hours between 5:30 PM and 6:30 PM and stored in RNAlater (Ambion) at 4°C until extraction. Total RNA was isolated from one retina per fish using RNeasy Lipid/Tissue Mini Kit (Qiagen) according to the manufacturer’s instructions. Samples were homogenized in 700 µL of Qiazol (Qiagen) containing 1% of β-mercaptoethanol (Sigma) with a rotor stator homogenizer (VMR) and put through an “on column” DNAse digestion using DNAse I (Qiagen). RNA concentration was determined by a Nanodrop spectrophotometer (GE Healthcare 28 9244-02) and Agilent 2100 Bioanalyzer (Agilent RNA 6000 NanoChip). For each sample, 100ng of total RNA was reverse transcribed using qScript cDNA Supermix (Quanta Biosciences) as per manufacturer’s instructions. cDNA was diluted 1:10 in Nuclease-free H_2_O (Ambion) and stored at −20°C until use. Primers were designed using the Primer 3 algorithm in Geneious R.9.1.7 [57]) to amplify cDNA products. All primers were validated using a standard serial dilution to determine efficiency under MIQE guidelines (Bustin *et al*., 2009). Dissociation curves were analyzed in 7500 Software v1.4.1 (Applied Biosystems, 2011) and only single products were detected. *β-actin* was used as endogenous control gene for normalization (Tang *et al*., 2007; Fleisch *et al*., 2013). Primer sequences can be found in STAR Methods. Each qRT-PCR reaction consisted of 2.5 μL of 3.2 μM primer solution, 5 μL of 2x (*Dynamite*) qPCR MasterMix (MBSU, University of Alberta) and 2.5 μL of cDNA in a 10 μL reaction volume. Technical replicates were performed in triplicate. RT-qPCR reactions were run on 7500 Fast Mode (pre-incubation 95°C, 2:00 min; 2 step amplification 95 °C 15s, 60 °C 1:00 min; 40 cycles; dissociation 95°C 15s, 60°C 20s, 95°C 15s, 60°C 15s) using the 7500 Fast Real-Time PCR System (Applied Biosystems). Biological samples (n= 5-6) each represent retinas from independent fish.

### Western Blots using custom anti-Nrl antibody

Adult zebrafish neural retinas were dissected and total protein was extracted by homogenizing samples in a standard lysis buffer (4mL 0.5M Hepes, 20mL 100% glycerol, 10mL 5M NaCl, 0.039g MgCl_2_, 40uL 0.5M EDTA, 100uL Triton X-100, 65.86mL ddH_*2*_O) with 1:200 protease inhibitor cocktail (EMD Millipore/VWR catalogue #CA80053-852, Darmstadt, Germany) with a rotor stator homogenizer (VWR catalogue #47747-370, Radnor, PA, USA). Homogenate was centrifuged at 13000 rpm for 8 minutes and supernatant was collected. Total protein in the supernatant was quantified using a Qubit Fluorometer (Invitrogen calatogue #Q32857, Carlsbad, CA USA). 30ug of protein was diluted with 2X SDS loading dye (12.5 mM Tris, 2% glycerol, 0.4% SDS, 0.2% β-mercaptoethanol, 0.1% bromophenol blue), and loaded into a 10% acrylamide stacking gel, followed by a 25% acrylamide separating gel. Protein was then transferred to a nitrocellulose membrane and blocked with 5% skim milk in tris buffered saline solution with 0.1% Tween (TBST). Membranes were probed with a custom polyclonal antibody raised against zebrafish Nrl, diluted 1:2000 in TBS. Antigen for the custom anti-Nrl antibody (commissioned from Genscript, Piscataway, NJ, USA) was recombinant protein matching the C-terminal 112 amino acids of zebrafish Nrl (residues 301-412 of GenBank: AAI63220.1). Antibody was affinity purified from the serum of a rabbit host. The blots probed with α-Nrl antibody were detected using 1:5000 goat α-rabbit HRP secondary antibody (Jackson Immunoresearch catalogue #111-005-003, West Grove, PA, USA) in TBST with 1% milk. Blots were developed using SuperSignal West Fempto Chemiluminescent substrate (Thermo Fischer scienctific catalogue #34095. Waltham, MA, USA) and visualized using ChemiDoc MP Imaging System (Hercules, CA, USA. Catalogue #17001402). The blot was then stripped and re-probed with a 1:5000 dilution of anti-β-Actin antibody (Sigma catalogue #A2066, St. Louis, MO, USA) in TBST with 1% milk. The anti-β-Actin antibody was detected using a 1:10000 dilution of goat α-mouse HRP secondary antibody (Jackson Immunoresearch catalogue #115-035-003, West Grove, PA, USA). The intensity of the bands was then analyzed using ImageJ (National Institutes of Health, Bethesda, MD, USA), to calculate the ratio of the intensity of the Nrl immunoreactivity compared to the intensity of the β-actin bands.

### Wholemount immunostaining

Anaesthetized larvae were fixed with 4% paraformaldehyde in 0.1M phosphate buffer with 5% sucrose pH 7.4; fixation was at room temperature for at least two hours, or overnight at 4°C. Fixed larvae were washed from PFA with phosphate buffered saline pH 7.4 with 0.1% Tween20 (“PBSTw”), then prepared for immunohistochemistry. Fish were washed in pure water 5 minutes, −20°C acetone for 7 minutes, rinsed out of acetone with PBSTw + 1.0% DMSO, and then blocked for at least 30 minutes in 10% normal goat serum (ThermoFisher) in PBSTw. Blocking solution was drained away, and primary antibody was applied. Antibody incubations were mixed 1:100 for 10C9.1 or 1:500 for other primary antibodies, and 1:1000 for secondary antibodies, and incubated at 4°C at least overnight. Larvae were washed from antibody over 2×5min and then 2×1hr washes in PBSTw, and then transferred to 70% glycerol in PBSTw until equilibrated. The retinas of equilibrated larvae were then prepared for mounting.

### Wholemount larval retinal dissections

For mounting whole retinas, glycerol-equilibrated larvae were relieved of their lenses using microscalpels (electrolytically-sharpened tungsten wire needles) as previously described [58]. After de-lensing, the corneal and scleral tissues covering the retina were cut and folded back, and the retina removed from the socket with the tungsten needles using a scooping manoeuvre. The retinas were then positioned vitreal-side down upon slides, and cover slips positioned upon the scleral side. Excess 70% glycerol was added as a mounting medium, and the slides were then imaged as described below.

### Cryosectioning and immunocytochemistry

Larvae were fixed as described above and, after PFA was washed away, subjected to a graded series of sucrose washes, from 5% to 20% sucrose in 0.1M phosphate buffer, and cryoprotected overnight in the final 20% sucrose stage. After, the fixed larvae were transferred to a 2:1 mixture of 20% sucrose buffer : OCT cryosection fluid (TissueTek) for one hour, then transferred to a final 1:1 mixture of the same. After equilibrating to this new buffer, fixed larvae were embedded into plastic molds, frozen at −80°C for at least two hours, and then sectioned at 10 um thickness. Cut sections were allowed to air dry 30 minutes before storage at −80°C at least overnight. Retrieved sections were warmed to room temperature over 20 minutes, then washed 3×5min in PBS + 0.1% Tween20 (“PBSTw”) to remove sectioning media residue. Sections were then blocked and stained as for wholemount immunohistochemistry (above), then covered with 70% glycerol in PBSTw as a mounting medium, mounted under a coverslip, and imaged as previously described [59].

### *in situ* hybridization on retinal cryosections

*In* situ hybridization on frozen sections was performed as previously described [38]. Briefly, frozen sections were thawed and rehydrated, then immediately re-fixed in 4% PFA to help tissue adhere to slides. Sectioned tissue was digested briefly with proteinase K and then re-fixed with 4% PFA. Sections were then acetylated with a mixture of triethanolamine and acetic anhydride, then dehydrated in a graded ethanol series with diluted 2x sodium citrate buffer. Tissues were pre-hybridized in Hauptmann’s buffer, then hybridized with riboprobe overnight at 70°C using 1µg/mL DIG-labeled riboprobe. Probe was washed off in a graded series of sodium citrate buffer diluted in maleate buffer as previously described. Sections were blocked and then incubated with anti-DIG-conjugated alkaline phosphatase antibody (Roche) at 1:5000 dilution overnight. Alkaline phosphatase chromogen reaction was performed, terminated with excess alkaline phosphate buffer and subsequent fixation with 4% PFA. Sections were mounted with glycerol. Developed sections were imaged on an Axioscope A.1 microscope (Carl Zeiss MicroImaging, Oberkochen) with 12 bit MacroFIRE camera (Optronics, Goleta, CA, USA).

Riboprobes to detect *rh1, sws1* and *rh2* opsins were produced as per our previous methods [36]. Riboprobes against *rh2* opsin a cocktail including DIG-labelled riboprobes against all of *rh2-1, rh2-1*, and *rh2-3*, akin to our previous methods [36].

Riboprobes to detect *nrl* and *nr2e3* were produced using primers listed in Table S1, cloned into pCS2+ or used directly as template to produce riboprobes of 972 and 867 bp, respectively.

### Confocal and stereomicroscopy

Confocal microscopy was performed with an LSM 700 confocal microscope mounted on a Zeiss Axio observer. Images were acquired with ZEN 2010 (v6.0, Carl Zeiss AG, Oberkochen, Germany). Micrographs were taken using 63x oil immersion (numerical aperture of 1.4) and 20x objectives (numerical aperture of 0.8). Images acquired with the confocal microscope were captured with gain adjusted to avoid any empty or saturated pixels, and after acquisition, image minima, maxima, and gamma were adjusted in Fiji (ImageJ, Version 2.0.0-rc-54/1.51h, NIH, Bethesda, MD, USA) to improve contrast. Stereoscopy images were taken as previously described [60] using brightfield and fluorescent channels in separate photos. Where relevant, brightfield images were converted to grayscale and merged in Fiji with fluorescent channel images to improve GFP visibility.

### Transmission Electron Microscopy

Adult eyes with the lenses removed were fixed overnight at 4°C. Fixative was 2.5% glutaraldehyde, 2% paraformaldehyde in 0.1M phosphate buffer PH 7.4. Fixative was washed out of the samples, which were then subjected to a graded series of ethanol washes, and then infiltrated with resin and embedded. Samples were then sectioned on a Richert-Jung Ultracut E Ultramicrotome to sections of 70-90 nm thickness. Gridded sections were then stained with uranyl acetate and lead citrate, and imaged at 80 kV on a FEI COMPANY transmission electron microscope, model Morgagni 268 (FEI company, Hillsboro, Oregon). Images acquired with a Gatan Orius CCD camera using Gatan DigitalMicrograph image acquisition software, version 1.81.78 (Gatan, Inc., Pleasanton, CA).

### Wholemount larvae photoreceptor quantification

In larval whole mounted eyes, the dorsal-ventral axis is readily apparent while imaging. After confocal imaging, using Fiji image analysis software a 100×100 µm boundary was positioned just dorsal to and centered above the optic nerve head of the retina. Within the delineated box, the *Process>Smooth* function of Fiji tamed wild background pixels, and then the *Process>Find Maxima* function was used to efficiently count fluorescently labeled photoreceptors.

### Quantification of adult nuclei in histological sections

To compare the relative abundance of retinal nuclei between adult wildtype and *nrl*^*-/-*^ mutant zebrafish, a counting region of interest (ROI) boundary was first established, consisting of 100µm measured along the outer plexiform layer in the indicated region. From this, an ROI boundary was drawn to encapsulate all the nuclei within this stretch, taking into account the bends in the retinal tissue. All nuclei were hand-counted except for the non-HC INL nuclei, which were counted as for wholemount larval photoreceptors (above).

### Ultrastructural synapse and chromatin appearance quantification

For all quantification and analysis, transmission electron micrographs of synapses and nuclei were imported into Fiji, and analysed as described. To quantify synaptic vesicle density, a region of interest (ROI) boundary was drawn around a photoreceptor synapse that excluded bipolar and horizontal cell processes. The interior of the ROI was then processed in the following way in Fiji: First, *Process>Subtract Background*, rolling ball radius = 5.0 pixels, “light background”. Next, background pixel noise was suppressed using *Process>Filter>Median*, radius = 1.0 pixel. The image was then thresholded with *Image>Adjust>Threshold*, auto. Finally, *Analyze>Analyze Particles* was used to count particles, with minimum and maximum particle radius set to 0.001 and 0.005µm (based on previous calibration). The area of the total ROI was recorded as the area of the synapse.

To measure synaptic ribbon length, only ribbons meeting particualr criteria were included: only ribbons with a clear synaptic ribbon “head” were measured; numerous synapses with oblique cut angles showed shadows of synapse ribbons slightly out of plane with the image, and the lengths of these shadows varied considerably (data not shown). Thus, only synaptic ribbons cut *en face* were measured. The segmented line tool of Fiji was used to trace the length of each ribbon, calibrated to the scale bar in each micrograph.

## Statistical Analysis

Statistical analyses (described in associated Figure legends, including Mann-Whitney-U, Wilcoxon ranked sum tests) and plot generation for larval photoreceptor abundances, synaptic vesicle density and ribbon length, relative abundances of nuclei in retinal sections, and rod nuclear sizes were performed in R (version 3.4.1, R Foundation for Statistical Computing, Vienna, Austria).

All RT-qPCR data is presented as mean ± Standard Error of Mean (SEM) and is standardized to wild type (AB strain) transcript abundances. To analyze differences in mean expression between wild type and mutant larvae and retinal tissue, a Mann-Whitney Test was performed using GraphPad Prism (Version 7.02 for Windows, or 8.4.2 for Mac, GraphPad Software, La Jolla California USA, www.graphpad.com). Two outlier data points were removed, after their identification with both ROUT analysis (Q = 5%) (α=0.05) and Grubb’s test (α=0.05) in GraphPad Prism; the same two outliers were identified when was implemented (Q = 5%, α=0.05). Statistical significances are denoted in each respective Figure legend.

## Supplement to

**Figure S1.**
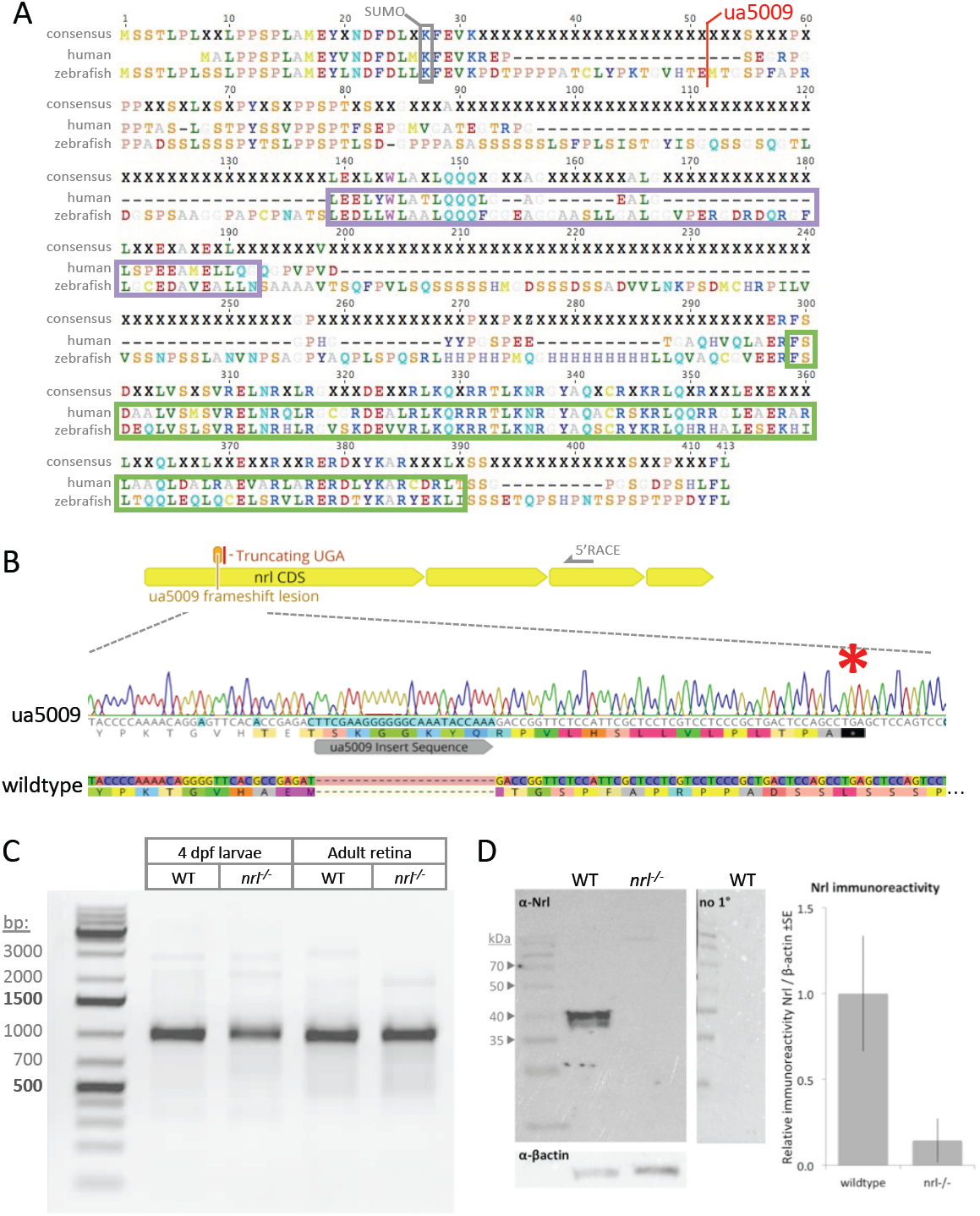
Most aspects of zebrafish Nrl protein are conserved with mouse and human Nrl, and the key protein domains are predicted to be absent in *nrl*^-/-^ frameshift mutants. Relates to Figure 1 and provides expanded views of data therein. **A**. Alignment to human Nrl reveals that the zebrafish Nrl homolog has insertions that make it relatively longer, but all key protein domains are recognizable. Maf N-terminal region (Maf_N) and basic leucine zipper DNA-binding (bZip) Domains are outlined and colour-coded purple and green, respectively as per Figure 1B. Both these domains are lost in *nrl*^-/-^ allele ua5009 where an insertion (see panel B) creates a frameshift and therefore abrogates normal translation of Nrl after residue 51. An experimentally-validated SUMOylation site at residue K20 of human or mouse Nrl is perfectly conserved in zebrafish (residue K27 and surrounding residues in zebrafish Nrl), and likely accounts for the doublet band of Nrl appearing on immunoblots of wildtype retina (panel D). **B**. CRISPR/Cas9 engineered *nrl* allele ua5009 is a frameshift (23 basepair insertion) near the beginning of the first coding exon of the *nrl* gene. The frameshift leads to a termination codon (*) shortly after the insertion. This truncation is predicted to eliminate the recognizable protein domains of Nrl (schematized in Fig. 1B and S1A) and is predicted to be a null allele. **C**. Characterizing the *nrl* transcript in *nrl*^-/-^ mutants shows no evidence of abnormal splicing, arguing against any cryptic exons being incorporated, and thus does not support any confounds to the prediction of a null allele. All *nrl* transcripts are amplified, regardless of their 5’ content, using 5’RACE (Random Amplification of 5’ cDNA Ends) with a primer positioned in the third coding exon (schematized in top right of panel A). Transcripts from wildtype (WT) and *nrl*^-/-^ mutant tissues showed no evidence of disparities between genotypes, in either larvae or adult retina. The identity (mutant vs. wildtype) of the transcripts was confirmed by sequencing. This is an expanded view of data in Fig. 1H. **D**. Nrl protein is lost in adult *nrl*^*-/-*^ mutant retina. Blots are an expanded view of data in Fig. 1G, and additionally demonstrate lack of signal development when primary antibody is excluded (right side). Histogram displays quantification of Nrl immunoreactivity relative to β-actin for n=3 individual fish of each genotype. Doublet band of immunoreactivity is reminiscent of blots of mammalian Nrl where mutagenesis has demonstrated this represents a post-translational SUMOylation; the SUMOylation site is perfectly conserved in zebrafish Nrl (Fig. S1A).

**Figure S2.**
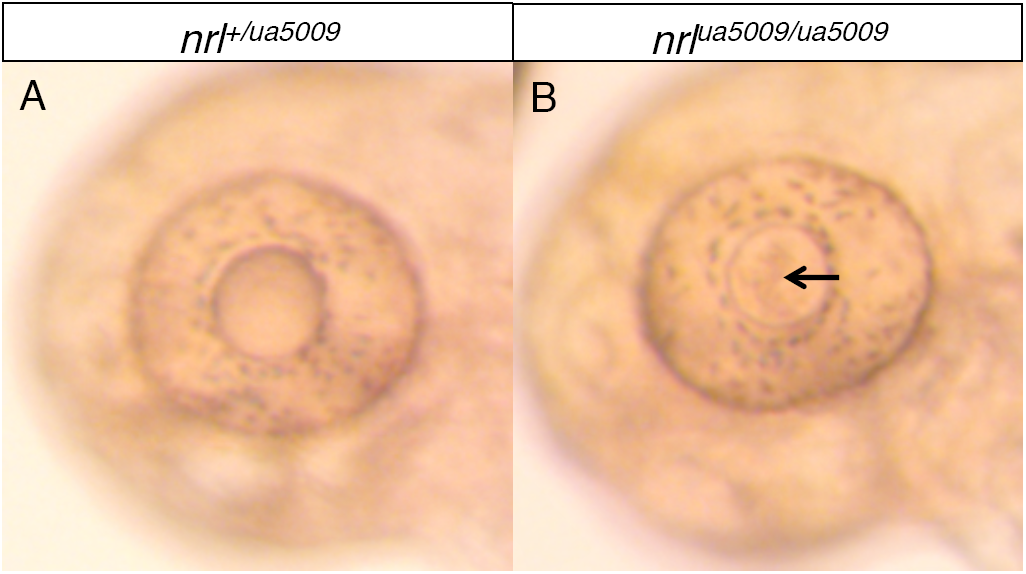
Proper zebrafish lens development requires *nrl*. Relates to Figure 4B. a small inclusion is visible in the lens of *nrl*^*-/-*^ larvae, and it remains detectable in adults (Fig. 4B’).

**Figure S3.**
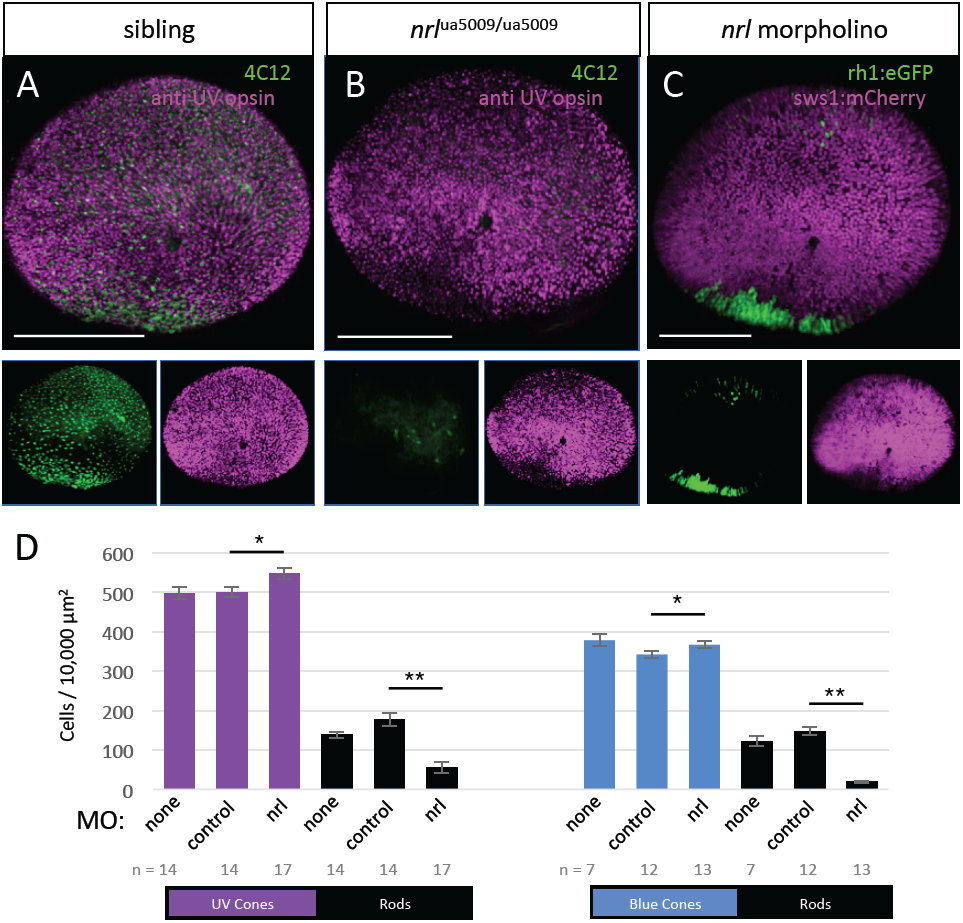
**Morpholino knockdown of *nrl* phenocopies the *nrl* mutation**, revealing drastic reduction of rod cells and increase in UV cones relative to wildtype. Wholemount retinas from 4dpf zebrafish, *en face* view (similar to Fig. 1C) with individual channels displayed below the merged channels. (A,B) green is 4C12 rod immunolabeling, while magenta is 10C9.1 UV cone immunolabeling; in panel C, rods express GFP from *Tg[rh1:GFP]* and UV cones express nfsb-mCherry (pseudocoloured magenta). **C**. 10ng of splice-blocking morpholino, targeting the first exon-intron boundary of *nrl* transcript, was injected into *nrl*^*+/+*^ zebrafish and sharply reduced the abundance of rod photoreceptors in the whole retina. **D. Rod and UV cone cell abundances change following *nrl* knockdown by morpholino**. Splice-blocking morpholino against *nrl*, or an equivalent amount of standard control morpholino, was injected into wildtype *nrl*^+/+^ zebrafish with the indicated fluorescent markers and rods and UV cones, or rods and Blue cones, were quantified within a 100 x 100 µm box positioned just dorsal to the optic nerve head. Rod abundance was substantially lower in nrl morphants relative to standard control-injected larvae, and a small (10%) but significant increase in UV cone abundance was observed. Morpholino injection decreased blue cone abundance, but less so with *nrl* morpholino compared to control morpholino. * is p<0.05, and ** is p<0.01 by Mann-Whitney U. n=number of individual larvae. Scale bars are 100 μm.

**Figure S4.**
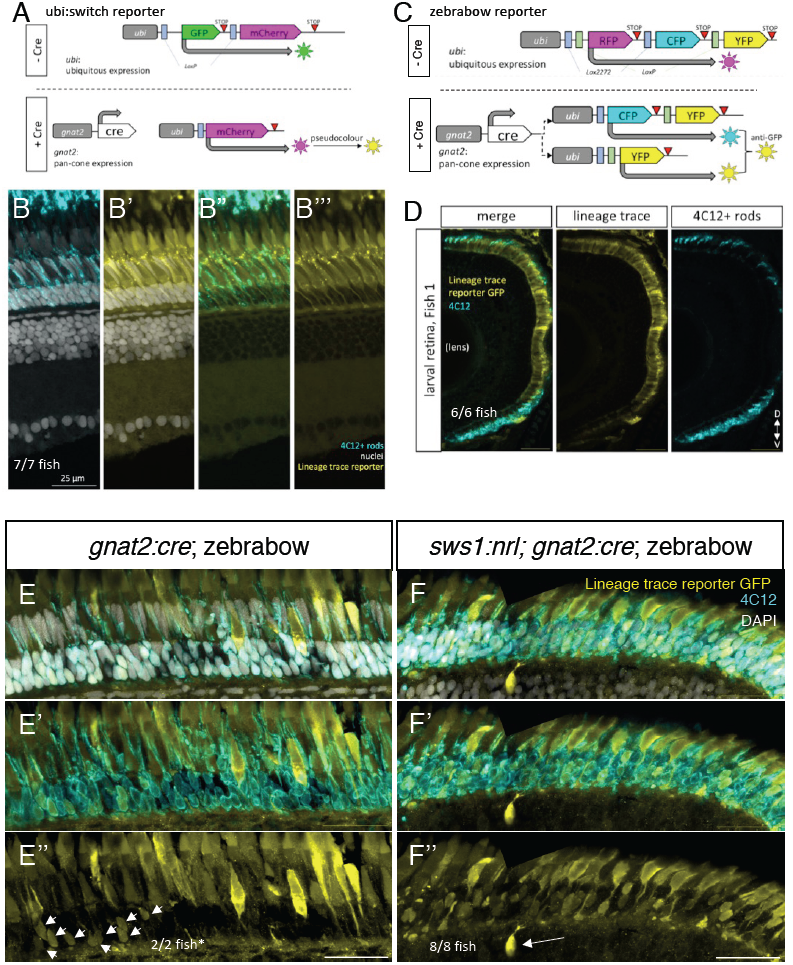
In wildtype zebrafish the cone photoreceptor lineage does not give rise to rod cells, though ectopic expression of Nrl shows the cone lineage has this capacity if artificially induced. Relates to genetically encoded lineage tracing in Figure 3C & D. **A**. Schematic of lineage tracing elements in ubi:switch transgenic zebrafish. In the absence of Cre recombinase, all cells in these fish express GFP (top). Cells of the cone lineage were engineered to express Cre recombinase (bottom, driven by promoter from the *gnat2* gene encoding cone-transducin-α) and thus all cells from the cone lineage permanently edit the reporter DNA to express mCherry protein (pseudocoloured to yellow). <u>In sum, all surviving cells of the cone lineage, and all their progeny, express mCherry (yellow) regardless of their subsequent cell fate</u>. **B**. The cone lineage in adult zebrafish retina gives rise only to cone photoreceptors, and no lineage-tracer-positive cells were found to co-localize the rod cell marker 4C12 in 7/7 animals. **C**. Schematic of zebrabow lineage tracing reporter is conceptually equivalent to panel A and gives rise to fluorescent protein (again pseudocoloured yellow) in all cells derived from the cone lineage. Differences to the ubi:switch reporter are that in the absence of Cre all cells of the body express red fluorescent protein (RFP) and the presence of Cre permanently switches cells to express a mixture of cyan and green fluorescent protein (CFP & YFP) that are detected with anti-GFP immunohistochemistry and pseudocoloured yellow. All cells of the cone lineage and their progeny fluoresce with reporter colour (yellow) regardless of their subsequent cell fate. **D**. In larval zebrafish the cone lineage has high fidelity towards generating only cone photoreceptors, and no rod cells (detected by immunomarker 4C12) co-localize the zebrabow cone lineage tracer. Results consistent in 6/6 larvae at 5 dpf (days post-fertilization). **E**. Adult wildtype (*nrl*^+/+^) fish bearing zebrabow cone lineage tracing broadly confirm results in panel B showing cone lineage labelling in 2/2 fish, *though in 1 of 2 examined retinas small clusters of lineage+ rods were observed; The largest such cluster (nine lineage reporter-positive rod cells, arrows) of this animal is shown, and 27 such cells were observed in five clusters, thus representing a tiny minority of rod cells and perhaps an artefact of stochastic recombination. Note typical wildtype (*nrl*^+/+^) retina in right side of E” containing no evidence of cone lineage trace reporter in rod cells. **F**. With transgenic expression of ectopic Nrl in UV cones via *Tg[sws1:Nrl]*, many rod cells (4C12+) now co-localize the cone lineage tracer. Thus UV cones expressing Nrl transmute to become rod cells. Also shown: a lineage reporter-positive bipolar cell (long arrow). There were a total of 3 bipolar cells labeled in this section, assessed between the CMZ and ONH.

**Figure S5.**
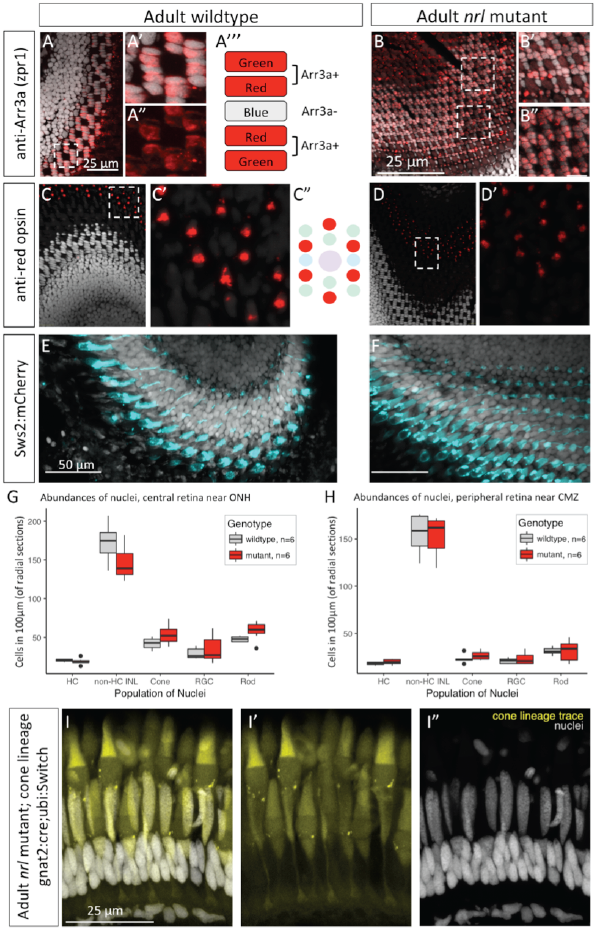
Adult retina of zebrafish *nrl*^-/-^ mutants have no overt phenotypes regarding cell type abundances or cone sub-types. Relates to Figure 4 regarding lack of expected photoreceptor phenotypes in adult *nrl*^-/-^ retina. **A-F**. Viewed in tangential (or somewhat oblique) cryosections, cone photoreceptors in adult zebrafish have a consistent and regularly repeated pattern (e.g. A” and C”) that allows sensitive detection of phenotypes or discrepancies, and none are detected in *nrl*^-/-^ mutant retina compared to wildtype. Sections are counterstained for cell nuclei (grey). Boxes outlined in main panels are included in adjacent insets (e.g. panel A has insets A’, A”, etc.). **A**,**B**. Double-cones (fused green- & red-sensitive cones) are detected as zpr1+ (an antibody against Arrestin3a) and display their typical repeating pattern, where each double cone flanks a zpr1-single cone (blue-sensitive cone). Double cones are zpr1+ and show typical inter-cellular organization/ positioning in *nrl*^-/-^ mutant retina. **C**,**D**. Red-sensitive cone opsin (*lws*) immunolabelling reveals red cones in their expected pattern amongst other cone types (schematized in C”) in both wildtype and *nrl*^-/-^ mutant retinas. **E**,**F**. Blue-sensitive cones, detected with transgene *Tg[sws2:mCherry]* that drives mCherry under the *sws2* opsin promoter (pseudocoloured cyan), shows the expected rows – a repeating pattern and regular spacing of blue-sensitive cones – in *nrl*^-/-^ mutant retina (F). **G**,**H**. Abundances of cell nuclei were determined in radial cryosections of adult retina (akin to sections in Fig. 3C & C’) by counting DAPI+ nuclei in each retinal layer. Radial sections selected for analysis were cut parallel to the nasal-temporal axis of the eye and transected the optic nerve head. Counts of various retinal cell nuclei were made from a region of retina delineated by measuring 100 µm along the outer plexiform layer and then counting the cells in each layer that were below or above this line. Counts were made on the basis of a single 0.58µm thick optical section, captured from a 10µm thick cryosection. Cell types are morphologically distinct and/or determined by their location in retinal layers: horizontal cell nuclei, HC; cells of the Inner Nuclear Layer other than horizontal cells, non-HC INL; cone cell nuclei, Cone; retinal ganglion cells, RGC; rod cell nuclei, Rod. Panel (G) quantifies cells in a central region, adjacent to the optic nerve head, whereas Panel (H) assesses peripheral retina near the ciliary marginal zone. Wilcoxon rank sum tests of each population of neurons by genotype revealed no significant differences after Bonferroni correction for multiple comparisons. n=6 individuals of each genotype. **I**. Lineage tracing of cone photoreceptors (from *Tg[gnat2:Cre]* and ubi:Switch reporter as per Fig. S1A,B) in adult *nrl*^-/-^ mutant retina shows normal abundance and morphology of cone cells. Further, there is no detectable signature of cone lineage tracing in the *nrl*^-/-^ rod photoreceptor cells.

**Figure S6.**
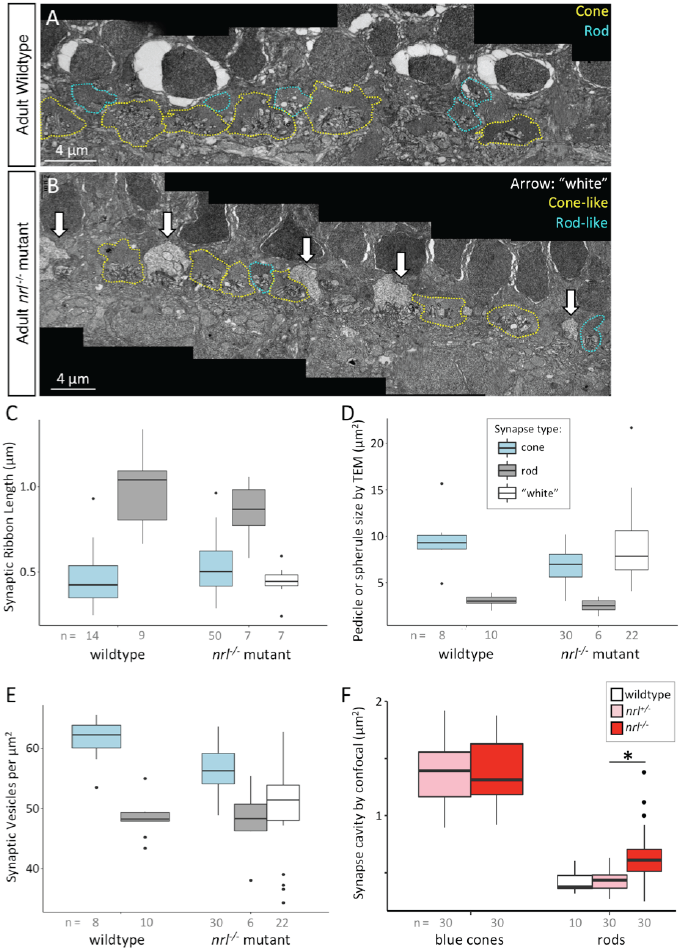
Photoreceptor synaptic terminals in adult retina of *nrl*^-/-^ mutant zebrafish suggest a requirement for Nrl in differentiation or maintenance of the rod synapse. Relates to Figure 6. **A**,**B**. Ultrastructure of Outer Plexiform Layer of adult retina, showing an expanded view of Fig. 6B,C. In wildtype adult zebrafish the photoreceptor synaptic terminals (panel A) include rod spherules (teal dotted lines) that are morphologically distinguishable from cone pedicles (yellow) based on their smaller size and smaller number of synaptic ribbons. In adult *nrl*^-/-^ mutant retina, the photoreceptor terminals (panel B) are disrupted compared to wildtype: a paucity of rod spherules are recognizable despite the presence of a normal abundance of rod cells. Further, only in adult *nrl*^-/-^ mutant outer nuclear layer, a subset of photoreceptor terminals were cone-like but extremely electon-lucent such that we denoted them ‘white synapses’ (white arrows). We suggest these white synapses may be *nrl*^-/-^ rod synapses because they are only present in mutants, and they would account for the disparity of only observing a paucity of rod spherules despite the large abundance of rod cells. Alternatively, the white synapses might be disrupted cone pedicles, but the data in panel (F) below is more consistent with *nrl*^-/-^ rod synapses being present and somewhat cone-like. **C-E**. Ultrastructural features of photoreceptor synaptic terminals were quantified in rod spherules, cone pedicles and ‘white synpases’ in each genotype. Rod and cones showed no striking difference based on genotype. White synapses are somewhat more cone-like (compared to a typical rod spherule) with respect to synaptic ribbon length (C), photoreceptor terminal size (D), and somewhat more variable with respect to density of synaptic vesicles (E). n= number of synaptic terminals, where wildtype data is from synapses imaged across two animals; mutant data from synapses imaged across 5 animals. Floating points are statistical outliers. **F**. Rod spherules were characterized in sibling vs. *nrl*^-/-^ mutant *Tg[rh1:gfp]* adult zebrafish by measuring the area of the synaptic cavity (the synaptic cleft contained between arms of GFP+ rod spherule) in confocal images of radial sections (e.g. Fig. 4D, D’). Similarly, the synaptic cavity of blue cone pedicles were quantified in fish bearing *Tg[sws2:mCherry]*. Blue cone pedicles were not apparently different between genotypes. Rod spherules had significantly larger synaptic cavities (and thus were somewhat more cone-like) in *nrl*^-/-^ rods (p = 4.319e-08, n = 10 synapses per fish, 3 fish per genotype for mutants and heterozygotes, and 1 wildtype fish), under Mann Whitney U comparison. Floating points are statistical outliers.

**Figure S7.**
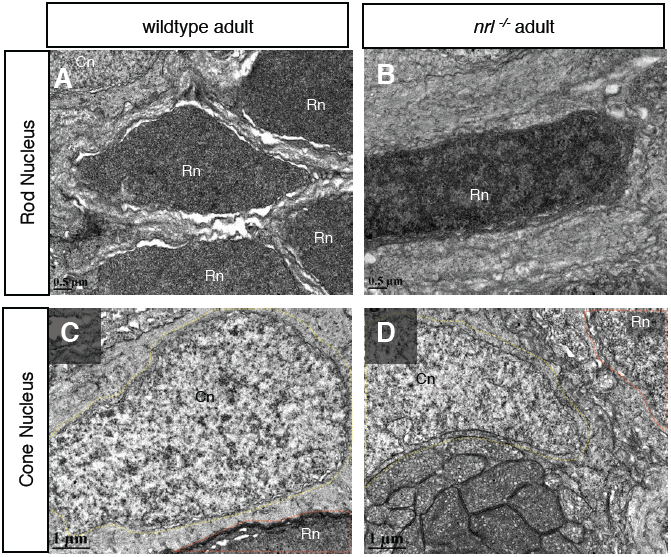
Photoreceptor nuclei in adult retina of *nrl*^-/-^ mutant zebrafish. In *nrl*^-/-^ adult retina the cone nuclei appear normal, but rod nuclei look somewhat ‘cone-like’, compared to wildtype, in that they have less homogeneous electron density.

**Supplemental Table S1.**
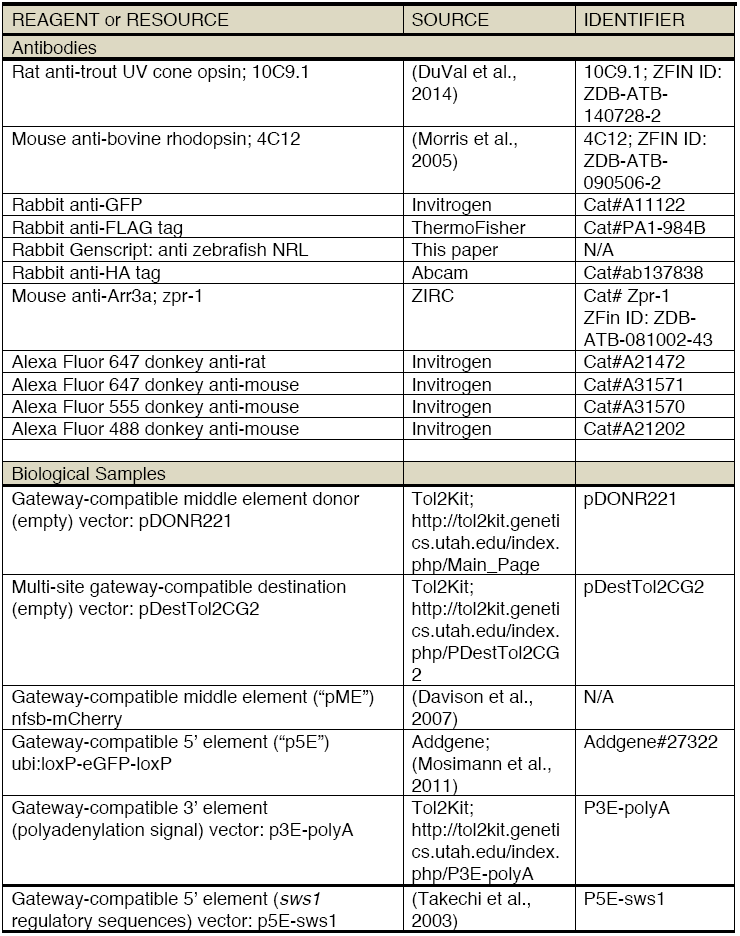
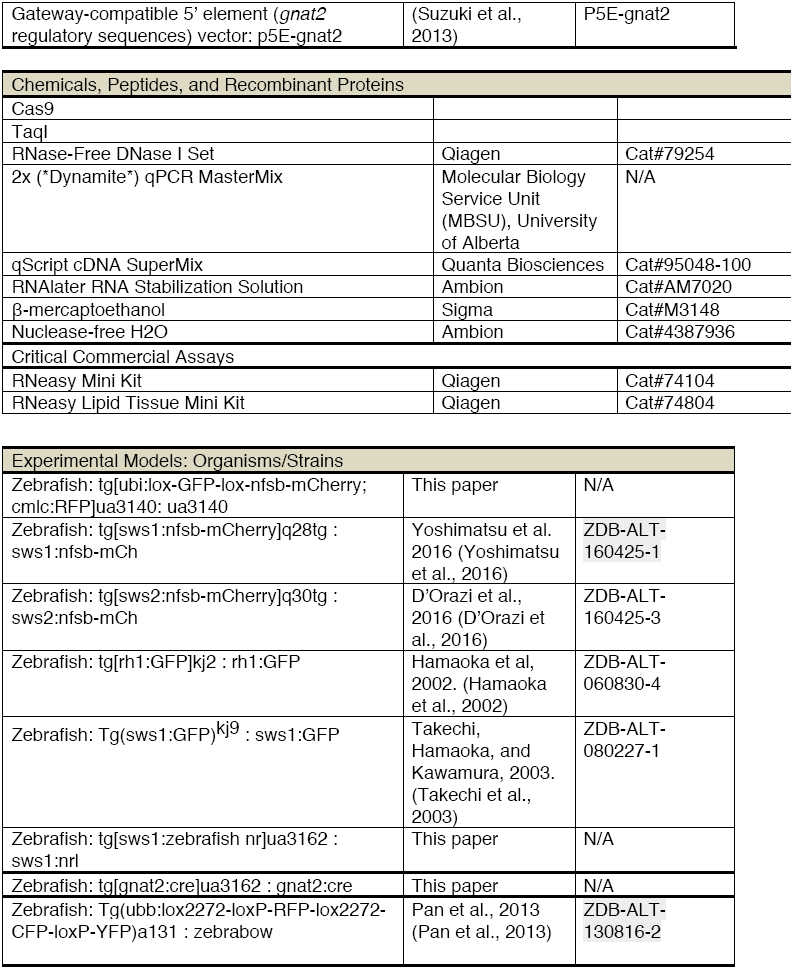
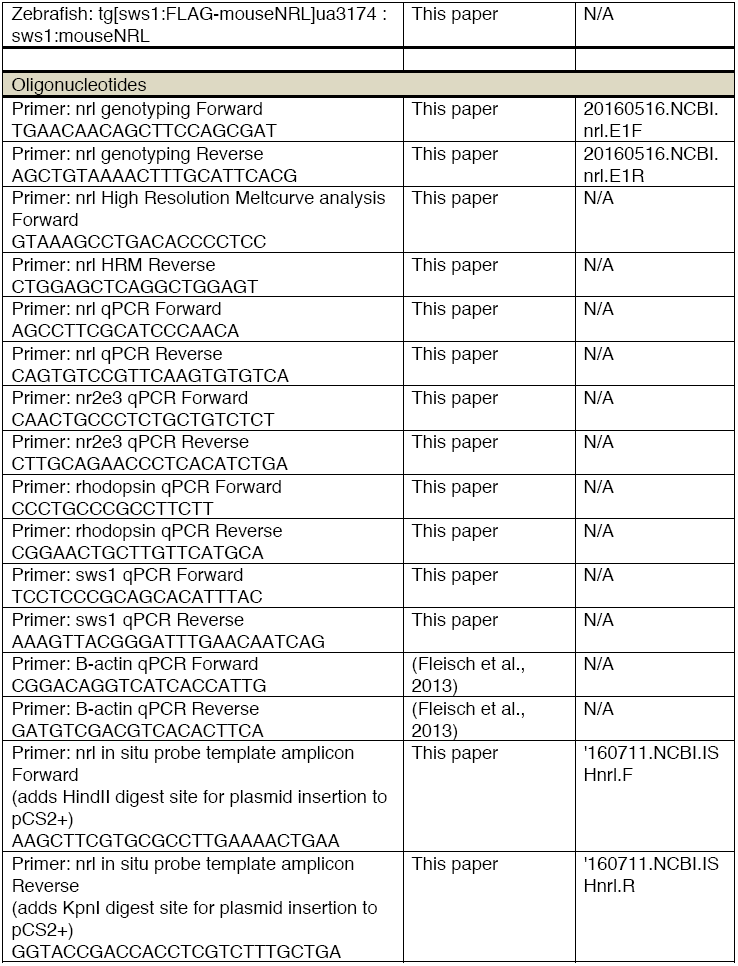
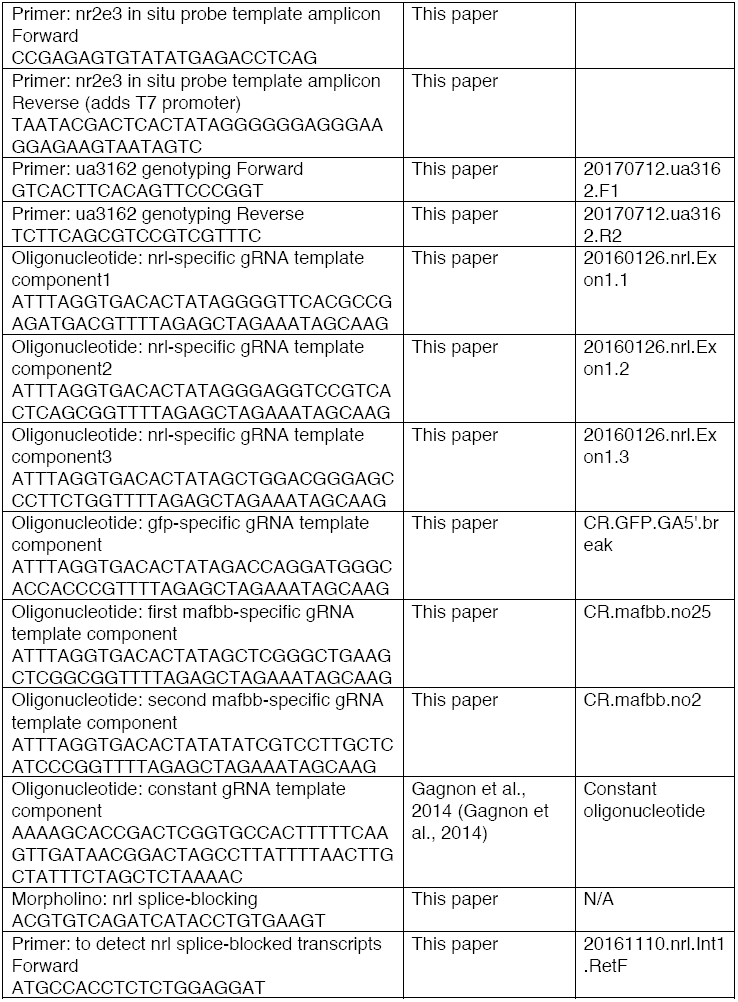
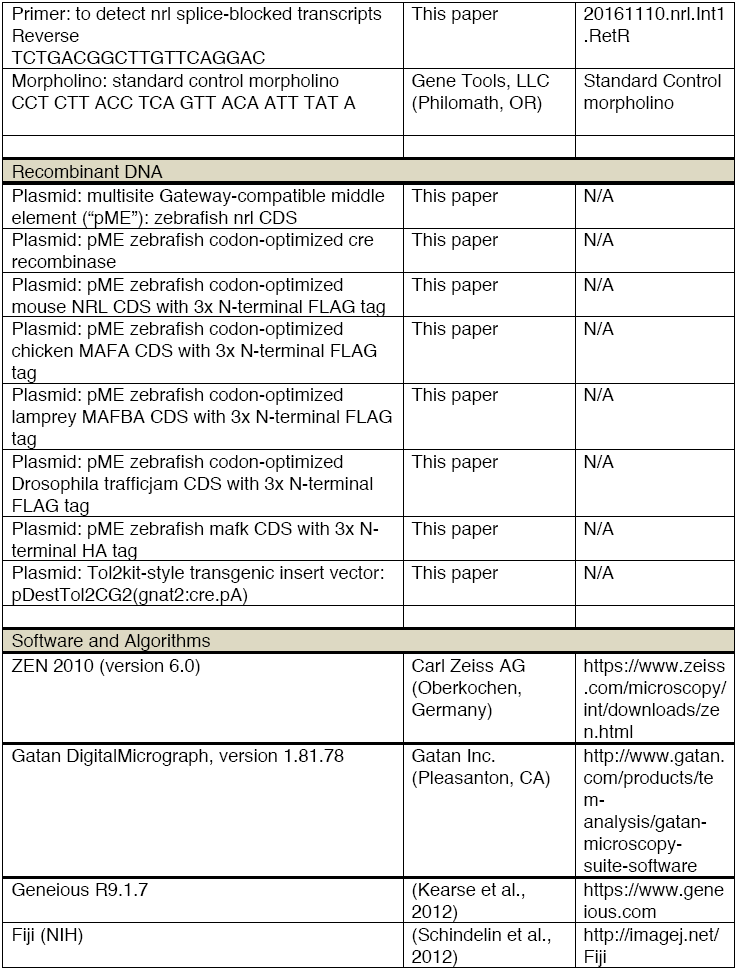
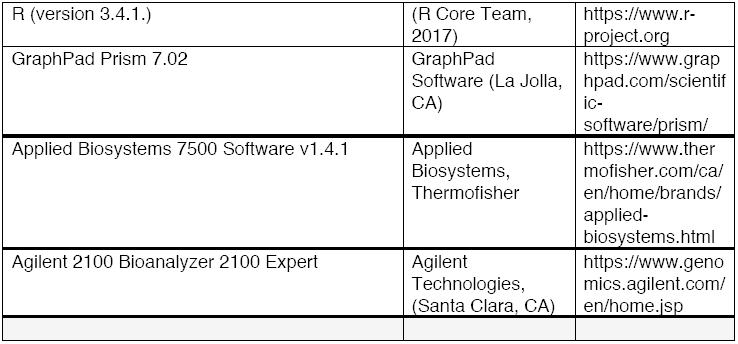
Key Resources

